# Growth/differentiation factor 15 controls number of ependymal and neural stem cells in the ventricular-subventricular zone

**DOI:** 10.1101/2022.12.02.518869

**Authors:** Katja Baur, Carmen Carrillo-García, Şeydanur Şan, Manja von Hahn, Jens Strelau, Gabriele Hölzl-Wenig, Claudia Mandl, Francesca Ciccolini

**Affiliations:** Department of Neurobiology, Interdisciplinary Center for Neurosciences (IZN), University of Heidelberg, Im Neuenheimer Feld 366, 69120 Heidelberg, Germany; Sorbonne University, 21 Rue de l’École de Médecine 75006 Paris, France. University of Heidelberg, Im Neuenheimer Feld 307, 69120 Heidelberg, Germany

**Keywords:** GDF15, GFRAL, EGFR, CXCR4, radial glia, subapical progenitors, neural stem cells, ganglionic eminence

## Abstract

Late in neural development, the expression of growth/differentiation factor (GDF) 15 increases in the germinal epithelium of the murine ganglionic eminence (GE), especially in progenitors with characteristics of neural stem cells (NSCs). However, the function of GDF15 in this region is still unknown. We here show that apical progenitors in the E18 GE also express the GDF15 receptor and that ablation of GDF15 promotes proliferation and cell cycle progression of apically and subapically dividing progenitors. A similar phenotype was also observed in the adult ventricular subventricular zone (V-SVZ). At both ages, increased proliferation leads to the transient generation of more neuronal progenitors, which is compensated by cell death, and to a permanent increase in the number of ependymal cells and apical NSCs. We also found that GDF15 receptor-expressing cells display immunoreactivity for the epidermal growth factor receptor (EGFR), which is also involved in progenitor proliferation, and that manipulation of GDF15 affects the expression of EGFR in mutant progenitors. Moreover, our data indicate that EGFR signalling in WT and mutant progenitors relies on distinct transduction modes. However, only exposure to exogenous GDF15, but not to EGF, normalized proliferation and the number of apical progenitors, indicating that alteration in EGFR signalling is not the main mechanism by which GDF15 affects proliferation in the embryonic GE.

Taken together, GDF15 directly regulates proliferation of apical progenitors in the developing GE, thereby affecting the number of total ependymal cells and NSCs in this region.

## Introduction

Before the onset of neurogenesis, the wall of the neural tube is a pseudostratified epithelium named ventricular zone (VZ) consisting of neuroepithelial cells (Huttner and Kosodo, 2005; McKay, 1997). Around the onset of neurogenesis, the latter start to express several glial-associated genes and transform into radial glia (RG) (Hartfuss et al., 2001; Levitt and Rakic, 1980; Misson et al., 1988; Rakic, 1972). All different types of neural cells derive directly or indirectly from RG through the generation of intermediate progenitors (Anthony et al., 2004; Bandler et al., 2017; Turrero García and Harwell, 2017). Apical intermediate progenitors derived from the RG in the ventral telencephalon are short neural progenitors (SNP) similar to those in the dorsal pallium (Tyler and Haydar, 2013) and subapical progenitors (SAP) (Pilz et al., 2013), which also display an apical attachment, but unlike apical RG and SNPs undergoing division at the apical border, SAPs divide between the apical and basal sides of the VZ. During development, neurogenesis occurs increasingly from progenitors located at the basal side of the niche, which also originate from apical RG. Basal progenitors undergo cell division at the basal side of the VZ, forming the secondary germinal epithelium termed subventricular zone (SVZ) (Haubensak et al., 2004; Kosodo and Huttner, 2009; Noctor et al., 2004; Petros et al., 2015; Pilz et al., 2013). They include asymmetric round neuronal progenitors with characteristics of transit amplifying cells and basal RG, which maintain a long basal process.

Progenitors in the GE contribute to the pool of adult NSCs in the adult ventricular-subventricular zone (V-SVZ) (Borrett et al., 2020; Young et al., 2007), which represents the largest neurogenic niche in the adult murine brain. In the V-SVZ, two morphological distinct types of NSCs coexist, which are referred to as apical and basal NSCs, as they display morphological characteristics of apical and basal RG (Baur et al., 2022). Whereas the origin of basal NSCs is still unclear, apical NSCs share a common apical RG progenitor with ependymal cells (Ortiz-Alvarez et al., 2019), whose proliferation and differentiation is affected by niche factors (Ferent et al., 2020).

In particular, from mid-development onwards the expression of the epidermal growth factor receptor (EGFR) and several potential ligands is increased in the tissue around the lateral ventricle, including in proliferating progenitors (Lazar and Blum, 1992; Zhang et al., 2023). Consistent with this, NSCs in the developing GE acquire responsiveness to EGF around E18 (Ciccolini, 2001; Ciccolini and Svendsen, 1998) and EGF represents the main mitogen for postnatal NSCs (Robson et al., 2018). However, despite its association with proliferation, pleiotropic effects have been observed downstream the EGFR activation including survival differentiation and migration (Abdi et al., 2018; Caric et al., 2001; Ciccolini et al., 2005; Scalabrino, 2022), which likely reflects different levels of EGFR activation (Burrows et al., 1997; Deribe et al., 2009) and differential recruitment of downstream pathways (Cochard et al., 2021). Notably, ablation of EGFR in developing neural precursors affects especially gliogenesis and astrocytic function (Robson et al., 2018; Zhang et al., 2023), likewise upregulation of EGFR activation in the postnatal brain promotes the acquisition of astrocytic characteristics in neural progenitors in the adult V-SVZ (Doetsch et al., 2002).

Growth/differentiation factor (GDF)15, a member of the TGF-β superfamily, is also expressed in the apical V-SVZ from late development onwards, however the function of the growth factor in this region is still unknown. In the adult brain, the choroid plexus epithelium of all ventricles represents the site of strongest and almost exclusive GDF15 expression in the adult CNS (Böttner et al., 1999; Schober et al., 2001). The neonatal brain displays GDF15 immunoreactivity not only in ependymal cells but also in the subependyma (Schober et al., 2001), suggesting that it may affect apical progenitor cells. In line with this, GDF15 has also been implicated in proliferation and invasiveness of human glioblastoma (Gritti et al., 1999; Shnaper et al., 2009; Strelau et al., 2008) and in the regulation of EGFR expression in hippocampal NSCs (Carrillo-Garcia et al., 2014). To investigate the potential role of GDF15 in the regulation of NSC behaviour, we have here analysed progenitors in the embryonic ganglionic eminence (GE) and in the postnatal V-SVZ of *Gdf15-*deficient mice. Our observations provide first evidence that GDF15 regulates the proliferation rate of apically and subapically dividing progenitors in the embryonic and postnatal VZ, thereby controlling the total number of ependymal cells and NSCs in the adult V-SVZ.

## Materials and Methods

### Animals and tissue dissection

All animal experiments were approved by the Regierungspräsidium in Karlsruhe, Germany. Time-mated pregnant (plug day =1.0) and young adult (8 weeks) C57BL/6 wild type (WT) (Charles River) and *Gdf15* knock-out/lacZ knock-in (Gdf15^-/-^) mice were killed by increasing CO_2_ concentration followed by neck dislocation. The dissection of the embryonic GE was done to maximise the excision of the germinal region as described before (Ciccolini and Svendsen, 2001) in ice-cold Euromed-N basal medium (Euroclone). After dissection, the tissue was dissociated mechanically. Dissection and dissociation of the SVZ were done as described before (Cesetti et al., 2009).

Gdf15^+/-^ matings were set up to obtain littermate embryos of the three genotypes (WT, Gdf15^-/-^ homozygous and Gdf15^+/-^ heterozygous). The generation of the Gdf15^-/-^ line and the genotyping have been already described (Strelau et al., 2009).

### BrdU labelling *in vivo*

Time-mated (E18) pregnant *Gdf15^+/-^* females from heterozygous matings were injected intraperitoneally with BrdU (10 μg/g body weight) once and sacrificed after 2 hours, or twice every two hours and killed 6 hours after the first injection.

### Primary cultures

Neurosphere cultures were established and induced to differentiate as previously described (Gakhar-Koppole et al., 2008). During proliferation human recombinant EGF (20 ng/ml), FGF-2 (10 ng/ml) (Peprotech) and GDF15 (10 ng/ml) (R&D) were added to the culture medium as described. Differentiating cells were fixed and processed for immunostaining at different time points after plating as indicated in the text. In some experiments BrdU (6.7 µg/ml) was added to the media for 16 hours to label dividing cells. Thereafter cultures were processed for immunocytochemistry.

Clonal cultures were obtained by plating one cell per well, either by means of limiting dilutions or FACS automated deposition, in 96 well/plates in the presence of the above-mentioned concentration of EGF and FGF-2 with or without GDF15 as detailed in the text. For secondary sphere formation, primary clones were mechanically dissociated and replated at a density of 1000 cells per well in 96 well plates in EGF and FGF-2 supplemented culture medium. Confluence was measured using the IncuCyte S3 Live-Cell Analysis Instrument (Sartorius).

### Fluorescence activated cell sorting (FACS)

Sorting was performed using a FACSVantage and a FACSAriaIII sorter (Becton Dickinson) as previously described (Carrillo-Garcia et al.; Cesetti et al., 2009; Ciccolini et al., 2005). For EGFR detection cells were stained in sort media with 20 ng/ml EGF conjugated to Alexa Fluor 488 or Alexa Fluor 647. For Prominin immunostaining cells were incubated one hour at 4°C with anti-Prominin-1 antibody conjugated to R-phycoerythrin (PE) or Brilliant Violet 421. Viable cells were revealed by propidium iodide (PI) exclusion (1 μg/ml, Sigma). All antibodies and labelled recombinant proteins used are listed in supplementary Table S1.

For β-galactosidase detection, dissociated cells obtained from Gdf15^+/-^ animals were incubated in a solution containing Fluorescein Digalactopyranoside (FDG, Molecular Probes) and labelled according to the manufacturer’s protocol.

### Whole mounts preparation and treatment

For whole mount preparations, brain tissue of 8-weeks-old animals or E18 embryos was dissected as described before (Khatri et al., 2014), and the GE or SVZ was directly fixed in a 3% PFA / 4% sucrose solution in PBS for 24 h. Alternatively, the tissue was incubated in a well of a 24-well-plate containing 1 ml Euromed-N (Euroclone) with 1x B27 supplement (Invitrogen) and as indicated either a solvent control, human recombinant GDF15 (10 ng/ml; R&D Systems, 9279-GD), human recombinant EGF (20 ng/ml; Peprotech, #AF-100-15), PD158780 (20 µM; Calbiochem/Merck, #513035) or AMD3100 octahydrochloride hydrate (6 µM; Sigma Aldrich, # A5602) at 37°C, 5% CO_2_ for 24h, and then fixed as described above.

For IdU labelling, E18 GEs were incubated in the same medium containing IdU (20 µM) for 1.5 h and then either directly fixed as above, or incubated a further 12 h in Euromed-N with B27 without IdU, followed by fixation. The fixed tissue was kept in PBS containing 0.01% sodium azide at 4°C until immunostaining.

### Western blot sample preparation

For western blot, the tissue of both GEs of one animal was mechanically dissociated in 600 µl Euromed-N with 1x B27 supplement, and the cell suspension was subsequently divided into 6 Eppendorf tubes á 100 µl. The cells were then treated with 100 µl additional medium containing, as indicated in the figure, either EGF (final concentration 20, 10, 5, 1 or 0.1 ng/ml) or GDF15 (final concentration 10 ng/ml) and incubated at 37°C, 5% CO_2_ for 7 minutes in the Eppendorf tube or overnight (O/N) in a well of a 48-well plate to ensure air flow. Thereafter, the cell suspensions were either spun down at 2000 rpm and resuspended in 50 µl ice-cold RIPA buffer containing 1x protease inhibitor, or spun down at 1500 rpm and resuspended in fresh pre-warmed medium. Cell suspensions were then further incubated at 37°C for 30 min, 1h, 2h or 4h as indicated in the figure and afterwards harvested as above. Samples were kept at −20°C until western blot.

Samples were mixed with 4x Laemmli buffer and run on a standard 7.5% SDS-polyacrylamide gel at a constant current of 30 mA per gel for 80-90 min on ice, then transferred onto a nitrocellulose membrane (Whatman Protran BA85, pore size 0.45 μm, Sigma-Aldrich) at a constant voltage of 20 V for 2h at RT. The membrane was blocked in 5% milk in tris-buffered saline containing 1% Tween-20 (TBS-T), and incubated with primary antibodies in TBS-T O/N at 4°C on a shaker. After washing with TBS-T, secondary antibodies conjugated to horseradish peroxidase were applied in 5% milk in TBS-T for 1h at RT on a shaker. Signal was detected with Amersham ECL Western Blotting Detection Reagent (GE Healthcare) using a Chemidoc Imaging System (Bio-Rad Laboratories).

All antibodies used for Western Blot are listed in supplementary Table S2.

### Immunofluorescence For cell cultures

Cells were fixed with 3% paraformaldehyde (PFA) in PBS containing 4% sucrose for 10 min at RT and permeabilised with NP-40 (0.5%). For BrdU stainings, cells were incubated with HCl 2 N for 30 minutes followed by a wash with sodium tetraborate 0.1 M pH 8.5. After permeabilisation, cells were incubated with primary antibodies O/N at 4°C and then with fluorescently labelled secondary antibodies for 1 hour at room temperature. The detection of O4 was done by incubating primary antibodies on live cells at 37°C. Nuclei were counterstained with 4’,6-Diamidine-2’-phenylindole-dihydrochloride (DAPI; Roche).

### For brain slices

Adult mice were anesthetised with an intraperitoneal injection of sodium-pentobarbital (2 ml/kg body weight; Narcoren®, Merial, Germany) and perfused with 4% paraformaldehyde (PFA), whereas brains from E18 WT and *Gdf15^+/-^*littermates were fixed by immersion in 4% PFA O/N. Thereafter, the tissue was cryoprotected by transfer into 30% sucrose O/N and then embedded in Tissue-Tek and frozen. Coronal slices 16 µm thick were cut with a cryostat (Leica CM3050S). Slices were permeabilised with 0.5% NP-40. For BrdU staining, slices were sequentially exposed to the following solutions in PBS: HCl 2 N, sodium tetraborate 0.1 M pH 8,5, glycine 0.1 mM, ammonium chloride 50 mM, and finally 5% FCS for one hour. Incubation with primary and secondary antibodies with DAPI was performed O/N at 4°C and for one hour at room temperature, respectively.

### For whole mounts

Whole SVZ / GE tissue was fixed as indicated above. When IdU was labelled, whole mounts were first incubated in 2N HCl at 37°C for 30 min; HCl was neutralized in 0.1 M Na-tetraborate, pH 8.5 for 30 min. All whole mount preparations were first briefly rinsed in 10 mM glycine and then permeabilized with 0.5% NP-40 for 10 min. To reduce background and unspecific staining, the tissue was incubated in 100 mM glycine and blocked with 5% FCS, 0.1% NP-40 for 1.5h at room temperature. Primary antibodies were applied in 0.1% NP-40 at 4°C O/N, and secondary antibodies and DAPI were applied in 5% FCS, 0.1% NP-40 for 2h at RT after washing. The tissue was mounted on glass slides using mowiol, with the apical surface of the GE/V-SVZ pressed flat against the glass coverslip. Both the β-catenin and fibroblast growth factor receptor 1 oncogene partner (FOP) primary antibodies are mouse monoclonal antibodies of the same immunoglobulin class. Therefore, for this double immunostaining both antigens were revealed using the same secondary antibody and each was distinguished based on the localization and morphology of the labelling, which is lining the cell boundaries or at the basal body of the cilia for β-catenin and FOP, respectively.

If not otherwise indicated, all solutions were diluted in PBS and all steps performed at RT. All antibodies used for immunofluorescence are listed in supplementary Table S3.

### Imaging of immunolabelled cells and slices

Fluorescent micrographs were acquired using an Axioplan 2 imaging microscope (Zeiss, Germany) equipped with a AxioCam digital camera (AxioCam HRc Zeiss, Germany) and AxioVision software (AxioVs40 V 4.5.0.00, Carl Zeiss Imaging, Germany). Confocal micrographs were acquired with a laser scanning confocal microscope (TCS SP2 scanning head and inverted DMIRBE microscope, 40x and 63x oil immersion HCX PL APO objective, Leica confocal scan (LCS) software; Leica, Germany).

### Confocal imaging of whole mounts

Whole mounts were imaged using a Leica TCS SP8 confocal microscope with a 40x or 63x oil immersion objective and LASX software (Leica). For the quantification, an average of three different regions of interest were chosen at fixed rostral, dorsal and ventral position of the GE or V-SVZ and averaged for the collection of a single data set. Images were acquired as z-stacks with 18-22 µm thickness with a z-step of 1 µm. Except for figure 2A, where single planes are shown, the images displayed in the figures are complete (when no reference to apical and subapical is made) or partial z-projections (5 z-planes for each apical and subapical) of the stack performed with Fiji/ImageJ (Schindelin et al., 2012).

### Image analysis

Images were analysed using Fiji/ImageJ software (Schindelin et al., 2012); Ki67, IdU/BrdU and phH3 cells were counted manually using the “Cell Counter” plugin if they contained visible nuclear staining.

Cells were considered dividing if the nuclei were labelled with Ki67 and the nuclear morphology showed a nuclear conformation characteristic of cells in meta-, ana-or telophase, in the DAPI and Ki67-channels. For the sake of clarity, throughout the study we use “dividing cells” only to refer to this way of detection, while cells positive for phH3 are exclusively termed “phH3^+^ cells”, as phH3 also labels cells in interphase and prophase, as well as late G2 phase. Mitosis was considered apical if the nucleus of the dividing cell was within two nuclei distance (∼ 10 µm) of the apical surface, and considered subapical if further away (as illustrated in Fig. 2A).

For each antigen at least three animals were analysed. For each mouse three sections at comparable rostrocaudal levels and separated from each other by five intervening sections (80 µm in total) were analysed. For fluorescence intensity measurements of EGFR, slices stained at the same time with the same antibody solutions, and imaged on the same day with constant confocal microscope settings (laser intensity, gain, pixel dwell time), were measured using Fiji/ImageJ. Raw immunofluorescence intensity was normalized by subtracting background fluorescence levels, i.e. fluorescence in cells considered negative for EGFR. To rule out any unspecific secondary antibody binding, fluorescence was compared to slices incubated with secondary, but not primary antibodies (2^nd^ only control); no difference was found between 2^nd^ only control and cells considered negative in EGFR-labelled samples, or between 2^nd^ only controls of different genotypes.

### Real time PCR

RNA isolation was performed using the TriFast reagent (peqlab) or Quick-RNA Micro Prep Kit (Zymo Research) according to the manufacturer’s protocol. RNA concentration was measured by optic densitometry using an Eppendorf BioPhotometer. Total RNA was retrotranscribed into cDNA using M-MLV reverse transcriptase with Oligo(dT)15 primers (Promega) according to manufacturer’s instructions. qPCR for the genes of interest was performed using the TaqMan gene expression assays and the StepOnePlus System (Applied Biosystems). Gene-specific TaqMan probes for the genes of interest *Gdf15*, *Gfral*, *Fgf-2*, *Fgf* receptors 1 and 2, and housekeeping genes 18s, *Gapdh* and *β-Actin* were obtained from Applied Biosystems as Assays-on-Demand (AOD) gene expression products. The AOD ID’s were *Gdf15* Mm00442228-m1; *Gfral* Mm02344882_m1; *Fgf-2* Mm00433287-m1; *Fgfr1* Mm00438923-m1; *Fgfr2* Mm00438941-m1; 18s, Hs99999901-s1; *Gapdh*, Mm00000015-s1; *β-Actin*, Mm00607939-s1. *Egfr* and *Gfral* transcripts as indicated were determined with PowerTrack SYBR Green (ThermoFisher) and the following primers: *Egfr* mouse forward 5’-ACCTCTCCCGGTCAGAGATG-3’, *Egfr* mouse reverse 5’-CTTGTGCCTTGGCAGACTTTC-3’ (Kefaloyianni et al., 2016); *Gfral* mouse forward 5’-GGGATGTTGGTTGGTGTCAG-3’, *Gfral* mouse reverse 5’-AGGCAGGTGTCTTCCATTGA-3’. Results were expressed as 2^−ΔΔCT^ which is an index of the relative amount of mRNA present in each sample.

## Results

### Expression of GDF15 receptor GFRAL in developing and adult apical progenitors

Previous studies have reported very low levels of *Gdf15* mRNA and protein in neural tissues, with the highest levels of GDF15 protein expression observed within the germinal epithelium lining the lateral ventricle and in the choroid plexus in neonatal and adult rats, respectively (Böttner et al., 1999; Schober et al., 2001). Consistent with these observations, we have previously shown that in the GE, *Gdf15* transcripts increase at late developmental age and remain high in the adult V-SVZ (Carrillo-Garcia et al., 2014). Moreover, we have found that *Gdf15* transcripts are expressed in neural progenitors displaying NSC potential in both the E18 hippocampus and GE and that in the latter region these NSCs also display high levels of the epidermal growth factor receptor (EGFR^h^) (Carrillo-Garcia et al., 2014), indicating that GDF15 may affect NSC development. Supporting this hypothesis, *Gdf15* transcripts were expressed in GE neurosphere-derived progenitors (supplementary Fig. S1A) and GDF15 immunoreactivity was present in cells expressing SOX2, a marker for NSCs (Ellis et al., 2004), and EGFR, which is upregulated in activated NSCs and cycling progenitors, in the adult V-SVZ (Fig 1A). The GDNF Family Receptor Alpha-Like (GFRAL), which has recently been identified as the GDF15 receptor (Li et al., 2017; Mullican et al., 2017; Yang et al., 2017), was also present in the E18 GE both when analysed at the level of RNA (supplementary Fig. S1B-D) and protein level (Fig. 1B, C). Notably, GFRAL was present especially in apical cells both in the embryonic GE (Fig. 1B) and in the adult V-SVZ (Fig. 1C) and often in cells which also displayed EGFR immunoreactivity. Taken together, our analyses show that GDF15 is particularly expressed in the germinal region of the GE and the adult V-SVZ. They also show that, like the growth factor, its receptor is also expressed in EGFR immunopositive progenitors.

**Figure 1:**
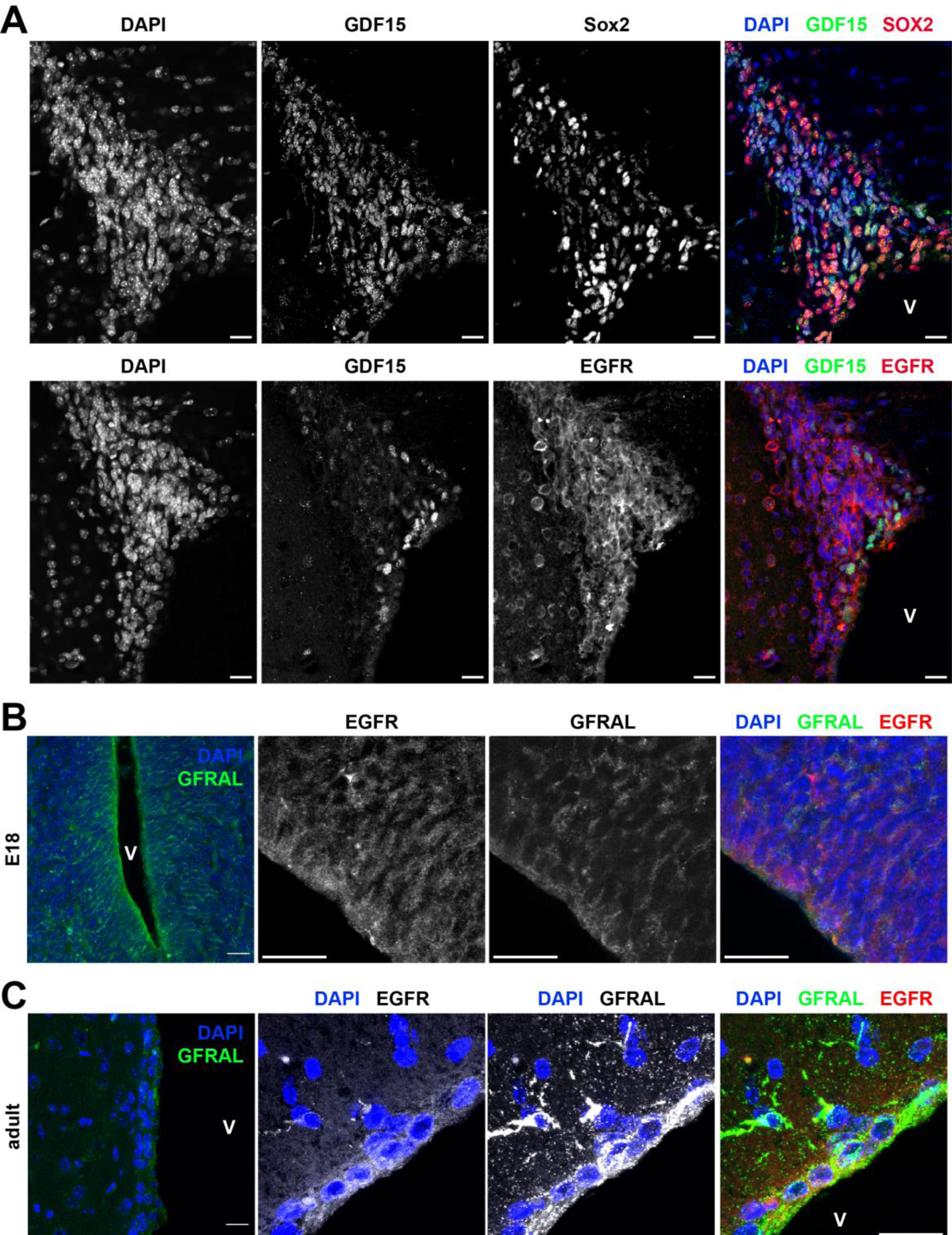
GDF15 and its receptor GFRAL are expressed in the V/SVZ. (A) representative confocal micrographs illustrating immunofluorescent staining of brain slices from 8w-old mice for GDF15 (green) and either Sox2 or EGFR (red). (B, C) Immunofluorescent staining of brain slices from E18 (B) or 8w-old mice (C) for GFRAL (green) in overview picture (left) and GFRAL (green) and EGFR (red) in a closeup image (right three images). DAPI (blue) was used as nuclear counterstain. V= ventricle. Scale bars = 20 µm.

**Table 1:** Extra BrdU incorporating cells exit the cell cycle in the *Gdf15^-/-^* GE after cell division. Quantitative analysis of the percentage of BrdU^+^ that are still cycling (BrdU^+^/Ki67^+^) or have exited the cell cycle (BrdU^+^/Ki67^-^) within the germinal (VZ/SVZ) and the differentiated region of the GE (GE) of E18 WT and *Gdf15^-/-^* embryos 6 hours after BrdU injection.

|  | WT |  | *Gdf15*<sup>-/-</sup> |  |
| --- | --- | --- | --- | --- |
|  | V-SVZ | GE | V-SVZ | GE |
| BrdU<sup>+</sup>/Ki67<sup>-</sup><br>(% BrdU<sup>+</sup>) | 23.9 ± 2.8 | 22.3 ± 1.0 | 42.5 ± 4.3** | 32.6 ± 3.6* |
| BrdU<sup>+</sup>/Ki67<sup>+</sup><br>(% BrdU<sup>+</sup>) | 76.0 ± 2.8 | 77.6 ± 1.5 | 57.4 ± 4.3** | 67.3 ± 3.0* |

### GDF15 affects the proliferation of apically and subapically dividing progenitors in the embryonic GE

Having observed that both GDF15 and its receptor are expressed in the GE and V-SVZ, we next investigated whether in this region, as in the hippocampus (Carrillo-Garcia et al., 2014), GDF15 also decreases progenitor proliferation. Quantitative analysis of the number of Ki67^+^ cells, showed that ablation of GDF15 led to a significant change in the number of cycling cells. However, unlike in the hippocampus, in the VZ of the embryonic GE we observed an increase in the number of cycling progenitors and in the number of dividing cells (Fig. 2A-C). The latter were scored based on the morphological nuclear appearance, see also the “Materials and Methods” section, which allowed us to detect dividing cells in anaphase and telophase (Fig. 2A white arrow). By distinguishing between progenitors undergoing cell division at the apical border, defined as the region lining the ventricle lumen with the thickness of around 10 µm, or at a subapical location, i.e., the region immediately adjacent the apical border along the apical-basal axis of the VZ (Fig. 2A), we were able to separately quantify apically dividing progenitors, i.e., RG and SNPs, and SAPs. We found that cell division was increased in both groups of progenitors in the mutant tissue at E18 (Fig. 2C). Notably, lack of GDF15 led to a similar effect in the adult V-SVZ with more cells undergoing apical division in mutant 8-week-old (8W) animals than in age-matched counterparts (supplementary Fig. S2A, B). To better understand the effect of GDF15 ablation on proliferation, we took advantage of whole tissue explants. After incubating the explants in medium containing IdU for 1.5 hours we quantified the number of cells undergoing DNA replication immediately thereafter or after further 12 hours of incubation in medium without IdU (Fig. 2D, E).

**Figure 2:**
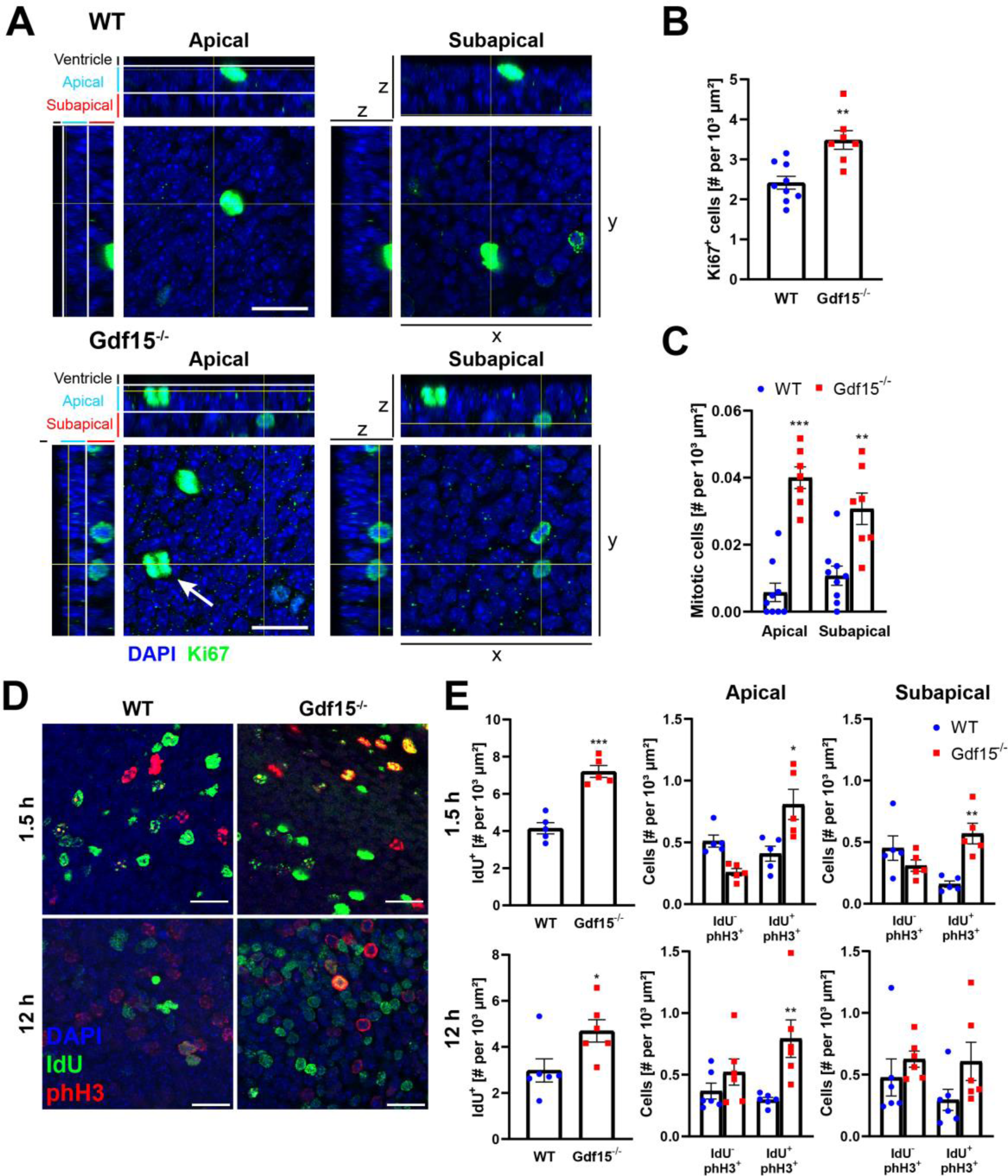
Gdf15^-/-^ mice display increased proliferation and number of proliferating cells in the VZ of the embryonic GE. (A) Immunofluorescent staining of E18 GE whole mounts for proliferation marker Ki67 (green). Large mages show single z-planes (x-y-axis) of apical and subapical regions; smaller images show orthogonally projected x- and y-planes (yz- and xz-axes, respectively, as indicated). The levels of ventricle, apical and subapical layer were defined as displayed on orthogonal projections. DAPI (blue) was used as nuclear counterstain. White arrow indicates a dividing cell. X, y, z = 3D axes. Scale bar = 20 µm. (B, C) Quantification of total number of Ki67^+^ cells (B), and dividing cells in the apical and subapical cell layers (C) in E18 whole mounts. (D) Immunofluorescent staining of E18 GE whole mounts for mitosis marker phH3 (red) and IdU (green). Images are taken at the apical side of the GE. The dissected tissue was incubated in IdU for 1.5h and directly fixed, or further incubated without IdU for 12h. DAPI was used as nuclear counterstain. Scale bars = 20 µm. (E) Quantification of cells labelled with IdU and phH3 directly after IdU application (1.5h) or after a 12h chase period (12h). Cells expressing phH3 were counted separately depending on location of the nucleus (apical or subapical) to infer the type of progenitor. Bars represent mean ± SEM; * indicate significance: *p<0.05, **p<0.01, ***p<0.001.

At both time points, more IdU^+^ progenitors were scored in the mutant tissue (Fig. 2E). Concomitant analysis of the localization of the nuclei displaying immunoreactivity to phospho-Histone H3 (phH3^+^) revealed that at the earliest time point, the extra IdU^+^ progenitors were progenitors undergoing mitosis both at the apical and subapical border (Fig. 2E). In contrast, the number of phH3^+^/IdU^-^ cells was not affected and it showed a trend decrease compared to the WT control, indicating a shortening of the time required for the transition between the S and M phases of the cell cycle and possibly of the duration of the M phase, in progenitors dividing in the absence of GDF15. Supporting this conclusion, the effect of the genotype on phH3^+^ cells, which include all cells in M phase, was smaller than the effect on Ki67^+^ dividing figures (Fig. 2 compare panels in C and E) which include only cells in advanced stages of mitosis. Finally, whereas at the earlier time point, the genotype did not affect IdU immunofluorescence levels (Fig. 2D, upper panels), 12 hours after IdU withdrawal, cells displaying light IdU immunoreactivity were present at the apical side (Fig. 2D, lower panels), which may represent second-generation subapical progenitors that have a 12-hour cell cycle, therefore significant shorter than other progenitors in the VZ of the embryonic GE (Pilz et al., 2013). Although these cells were not included in the quantification, they also appeared increased in the mutant tissue indicating that in the absence of GDF15 extra dividing cells in the VZ include also secondary progenitors. At the later time-point, only the number of apical phH3^+^/IdU^+^ cells was still increased in the mutant tissue, whereas mutant phH3^+^/IdU^+^ SAPs displayed only a trend increase that was not significant (Fig. 2E). We next analysed the percentage of cells which were still cycling after having replicated their DNA to investigate whether the increase in the number of proliferating progenitors in the mutant VZ is due to increased cell cycle re-entry (supplementary Fig. S2C, D). After 1.5 hours in the presence of IdU, compared to the WT counterpart, lack of GDF15 significantly altered the proportion of both mutant cycling and non-cycling IdU^+^ progenitors with a 3-fold and a 1.4-fold increase, respectively (supplementary Fig. S2C). Since cell cycle progression is faster in mutant than in WT progenitors, this indicates that more dividing progenitors re-enter the cell cycle in the mutant than in the WT tissue. However, after 12 hours, all the extra IdU^+^ cells in the mutant GE had exited the cell cycle (supplementary Fig. S2D), showing that the extra cycling cells in the mutant GE do not represent permanently cycling progenitors. We next investigated the cell division angles relative to an axis orthogonal to the ventricular surface, to see if apical extra proliferation is associated with an increase in symmetrically dividing cells (Supplementary Fig. S3A-E). Both at E18 and 8W, most apically dividing cells divided with a 0° angle (i.e., symmetrically) in both genotypes (Supplementary Fig. S3A-C). In the embryonic tissue, about 40% of dividing cells displayed a 45° angle (asymmetrically; Supplementary Fig. S3D), which is consistent with previous observations (Falk et al., 2017). This number decreased to less than 20% in the adult animals (Supplementary Fig. S3E), indicating a decrease in neurogenic divisions while NSC maintaining divisions persist (Chenn and McConnell, 1995; Zhang et al., 2004). Thus, the Gdf15^-/-^ V-SVZ has a higher number of cells undergoing mitosis but the distribution of division mode is not changed compared to the WT control at either age. Taken together, these data indicate that GDF15 affects cell division of neural progenitors also in the GE, as previously observed in the developing hippocampus (Carrillo-Garcia et al., 2014). However, in contrast to our previous observations, in the GE the absence of endogenous GDF15 leads to enhanced proliferation of VZ progenitors. Consistent with this, we found that compared to the WT counterpart, the number of apical Prominin-1-expressing (P^+^) progenitors was increased in the E18 mutant GE (Fig. 3A, B), which was reversed by exposing whole mount preparations of the E18 mutant GE to exogenous GDF15 for 24 hours (Fig. 3C). This treatment also normalized the number of cycling progenitors to WT levels (Fig. 3D, E), and it reduced the number of mitotic SAPs and apically dividing progenitors in explants of the mutant GE (Fig. 3F). In contrast, exposure to exogenous GDF15 largely did not alter the proliferation in the WT GE, with the exception of a significant increase in the number of apically dividing progenitors (Fig. 3F). Notably, the number of apically dividing progenitors in the WT and mutant tissue treated with GDF15 was virtually identical, indicating that the apparent opposite effect that exposure to GDF15 has on WT and mutant progenitors is likely due to the initial differences in the number of proliferating apical cells between the two genotypes. Since we have found that GFRAL is expressed in apical progenitors, our data indicate that GDF15 directly affects proliferation of VZ progenitors in the developing GE by decreasing proliferation of most apically and subapically dividing progenitors.

**Figure 3:**
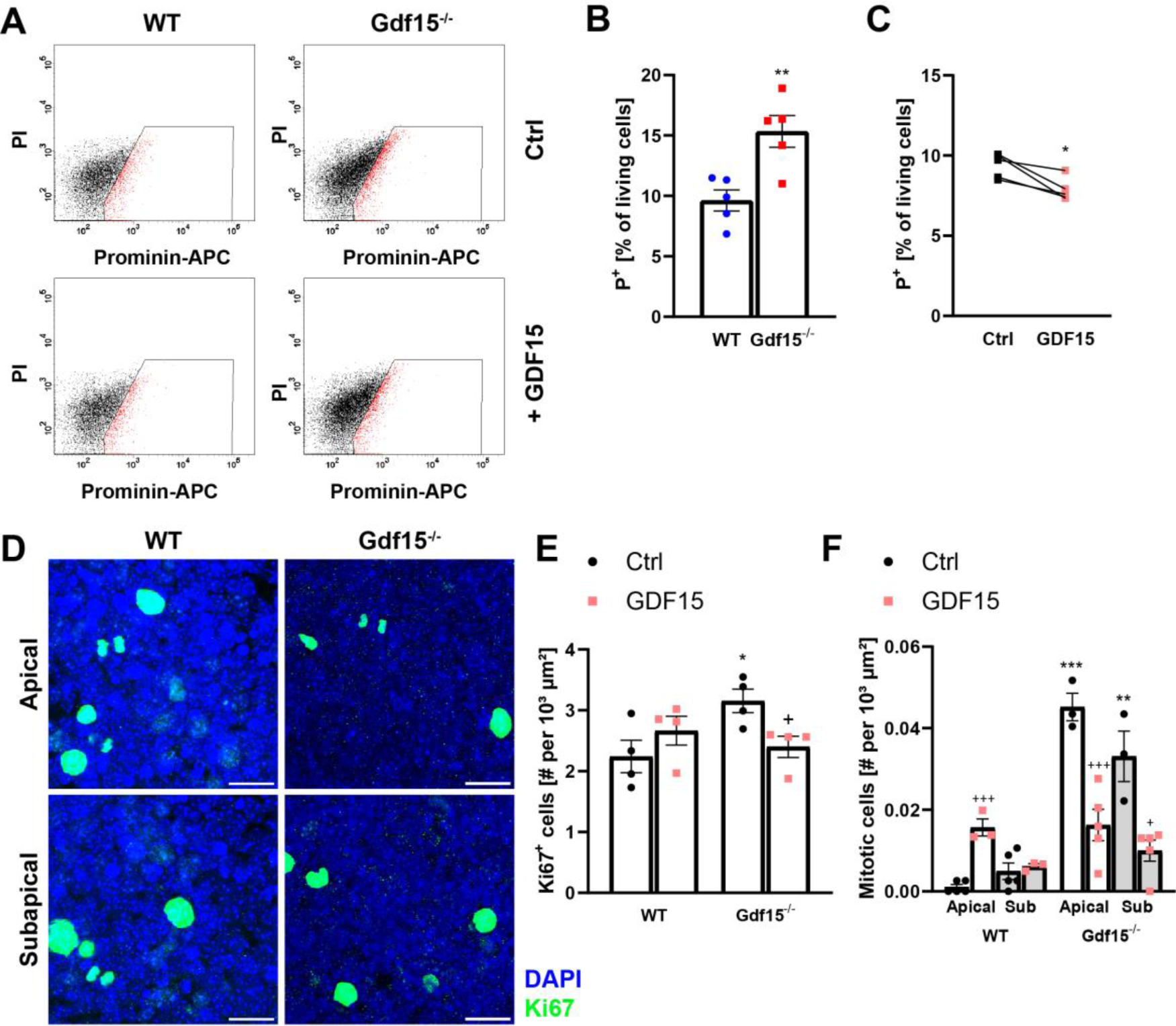
Application of exogenous GDF15 rescues phenotype. (A) Flow cytometry plots showing cells positive for Prominin-1 (P^+^, red) in the WT and Gdf15^-/-^ E18 GE, with GDF15 application (+ GDF15) and without (Ctrl). (B) Quantification of Prominin-positive cells (P^+^) as a percentage of total living (i.e., PI^-^) cells in Ctrl conditions. (C) Quantification of P^+^ cells after GDF15 treatment, as percentage of total P^+^ cells without treatment from the same animal. (D) Confocal images of E18 GEs treated with GDF15 overnight, with apical and subapical planes. Scale bars = 20 µm. (E, F) Quantification of Ki67^+^ cells (E) or dividing cells (F) in E18 GE whole mounts either incubated without (Ctrl) or with GDF15 overnight. Bars represent mean ± SEM; */^+^ indicate significance: */^+^p<0.05, **p<0.01, ***/^+++^p<0.001; * indicates significance to respective WT control, ^+^ indicates significance to respective untreated control.

### GDF15 regulates the proliferation of intermediate progenitors in the developing GE

We next investigated the nature of the extra proliferating progenitors in the VZ of the mutant GE. Cell cycle regulation affects neuronal output in the developing GE and in differentiating neurosphere cultures most immature neurons are generated from dividing progenitors (Ostenfeld and Svendsen, 2004; Suh et al., 2009). Moreover, our analysis of dividing cells indicates that the number of fast proliferating secondary SAPs are increased upon GDF15 mutation. Therefore, we next used differentiating neurosphere cultures to begin to characterize whether GDF15 mutation leads to the generation of extra neuronal progenitors. Consistent with our previous observations *in situ*, two days after induction of differentiation, Gdf15*^-/-^* neurosphere cultures contained significantly more cells replicating their DNA (Fig. 4A, B) and undergoing mitosis (Supplementary Fig. S4A, B) than their WT counterpart. This also resulted in a faster increase of Gdf15^-/-^ cell culture confluence within the first 24h of plating (Supplementary Fig. S4C). This difference could be prevented by adding exogenous GDF15 to the differentiation medium (Fig. 4B). Moreover, this difference was transient as it was not observed when cultures were examined at later stages of differentiation (Fig. 4C). No difference in the number of pycnotic nuclei was observed in cultures that had been differentiating for two days (Supplementary Table S4), showing that the change in cell proliferation observed in mutant cultures was not due to an effect of GDF15 on cell death. However, cell viability was compromised in mutant cultures that had been differentiating for 4 days in the absence but not in the presence of exogenous GDF15 (Supplementary Table S5). Besides proliferation and cell viability the genotype also affected neuronal differentiation. After 7 days in differentiating conditions, neurospheres originating from the E18 WT GE gave rise to significantly more TuJ1^+^ neurons than the Gdf15^-/-^ counterpart (Fig. 4D, E), whereas the number of O4^+^ cells was not affected by the genotype (Supplementary Fig. S4D). Finally, differences in the percentage of TuJ1^+^ neurons were not observed if cultures were allowed to differentiate for three further days (Fig. 4F) or if they were treated with exogenous GDF15. Taken together, these data indicate that GDF15, although it decreases the proliferation of neuronal progenitors, it promotes their differentiation and survival. We next investigated changes in proliferation and neurogenesis *in vivo* in deeper areas of the GE tissue, containing differentiated neurons. Analysis 2 and 6 hours after BrdU injection showed a significant increase in the number of dividing cells only at the latter time point (Fig. 5A), suggesting that they may represent more slowly proliferating neuronal precursors. Consistent with this, the number of cells expressing MASH-1, which in the developing GE is necessary both for the proliferation and specification of neuronal progenitors (Casarosa et al., 1999; Torii et al., 1999) was higher in dissociated cells obtained from the mutant E18 GE than in the tissue derived from their WT littermates (% immunopositive cells: WT = 22.84 ± 3.4; Gdf15^-/-^ = 30.51 ± 2.2; P<0.05). A similar change was observed in the postnatal SVZ, where progenitors expressing Ascl1 largely represent rapidly proliferating TAPs and pre-neuroblasts (Cesetti et al., 2009; Parras et al., 2004). In the postnatal ventral and mediolateral portion of the V-SVZ Ascl1^+^ progenitors were more abundant in Gdf15^-/-^ than in WT mice (Fig. 5B, C). Moreover, this increase was limited to the fraction of Ascl1^+^/Ki67^+^ proliferating progenitors (Fig. 5C) and was not observed at the level of the dorsolateral corner (Fig. 5D). A subset of Ascl1^+^ cells, but not pre-neuroblasts, also express Olig2, a helix-loop helix transcription factor associated to oligodendrocyte differentiation (Cesetti et al., 2009; Lu et al., 2002; Zhou and Anderson, 2002). We therefore next analysed the expression of Olig2 by immunohistofluorescence in the WT and Gdf15^-/-^ mediolateral V-SVZ. Independent of Ascl1 expression, the numbers of Olig2^+^ cells were not affected by the genotype (supplementary Fig. S5A). Likewise, we did not detect differences between total or proliferating DCX^+^/Ki67^+^ neuroblasts (Supplementary Fig. S5B, C) between the WT and mutant V-SVZ. We next investigated if cell death was also increased *in vivo* in the absence of GDF15, as this could explain why the extra-proliferating Ascl1^+^ progenitors in the Gdf15^-/-^ V-SVZ do not give rise to a higher number of postmitotic neuroblasts in the V-SVZ. Indeed, analysis by flow cytometry showed that, compared to the WT, the percentage of PI^+^ (i.e., dead) cells was higher in both populations in the mutant V-SVZ (EGFR^h^ cells: fold increase 2.00 ± 0.3382; *p* = 0.0369; EGFR^l^ cells: fold increase 1.380 ± 0.1094; *p* = 0.0511) and Annexin V positive cells were increased in mutant brain slices (Supplementary Fig. S5D). Taken together, our data indicate that both *in vitro* and *in vivo*, the increase in proliferation of mutant neuronal progenitors does not lead to increased neurogenesis, likely due to compensatory cell death.

**Figure 4:**
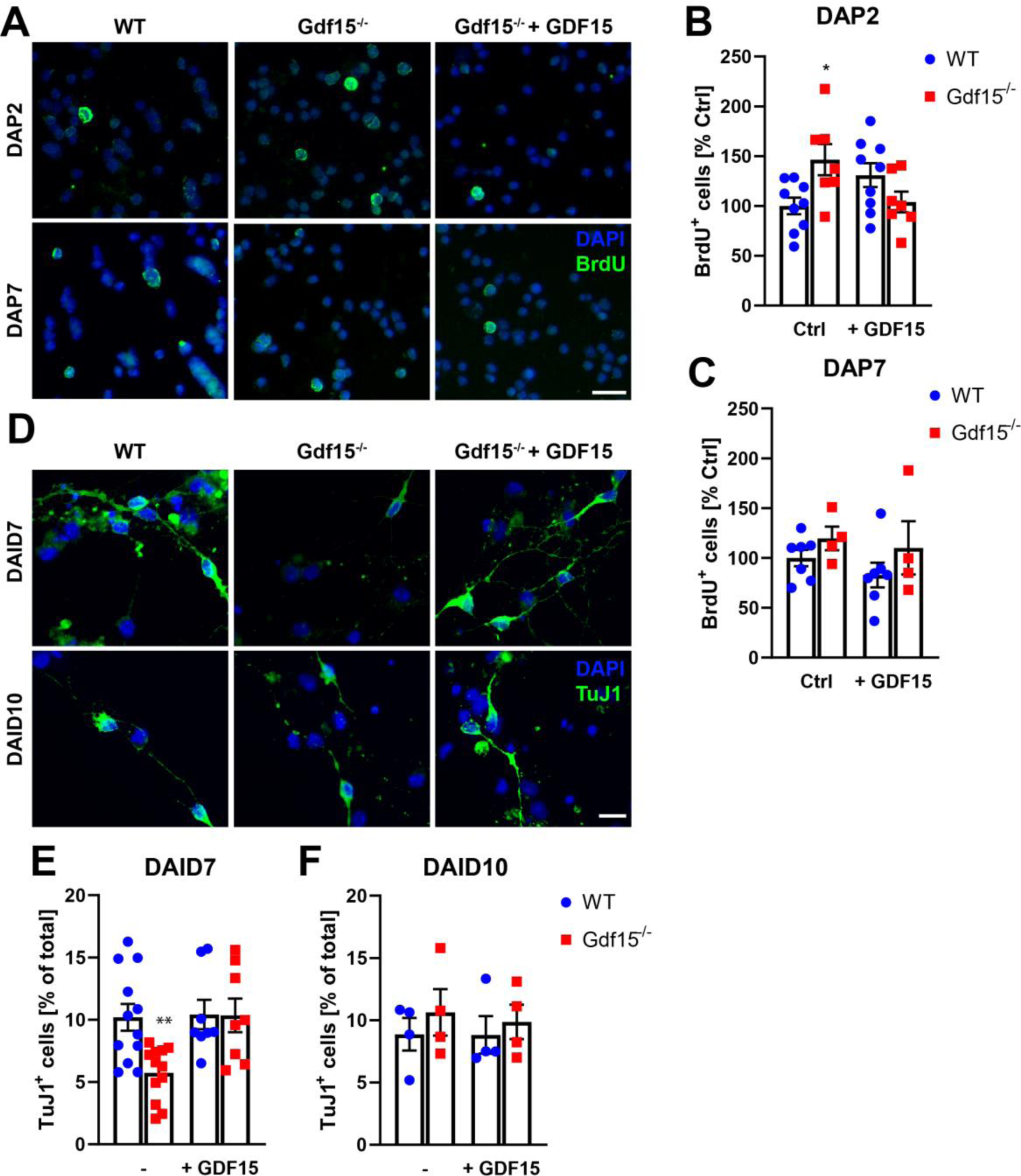
Primary Gdf15^-/-^ cell cultures show a transient change in proliferation and neurogenesis. (A) BrdU^+^ cells (green) and DAPI counterstaining of the nuclei (blue) in WT and Gdf15^-/-^ neurosphere cultures fixed two or seven days after plating (DAP2, DAP7) in differentiation medium. BrdU was added to the culture medium 24h before fixation. Scale bar = 20 µm. (B, C) Quantitative analysis of BrdU^+^ cells fixed two days (B) or seven days after induction of differentiation (C), with recombinant GDF15 added during differentiation (+GDF15) or without (Ctrl). (D) Representative examples of neurosphere cultures derived from the E18 GE fixed seven or ten days after the induction of differentiation (DAID7, DAID10) treated with exogenous GDF15 during the entire phase of differentiation (GDF15) or left untreated (Control) as indicated, stained for TuJ1. DAPI was used for nuclear counterstain. Scale bar = 20 µm. (E, F) Quantitative analyses of the number of TuJ1^+^ neurons in cultures fixed seven (E) and ten (F) days after the induction of differentiation. Bars represent mean ± SEM; * indicates significance: *p<0.05.

**Figure 5:**
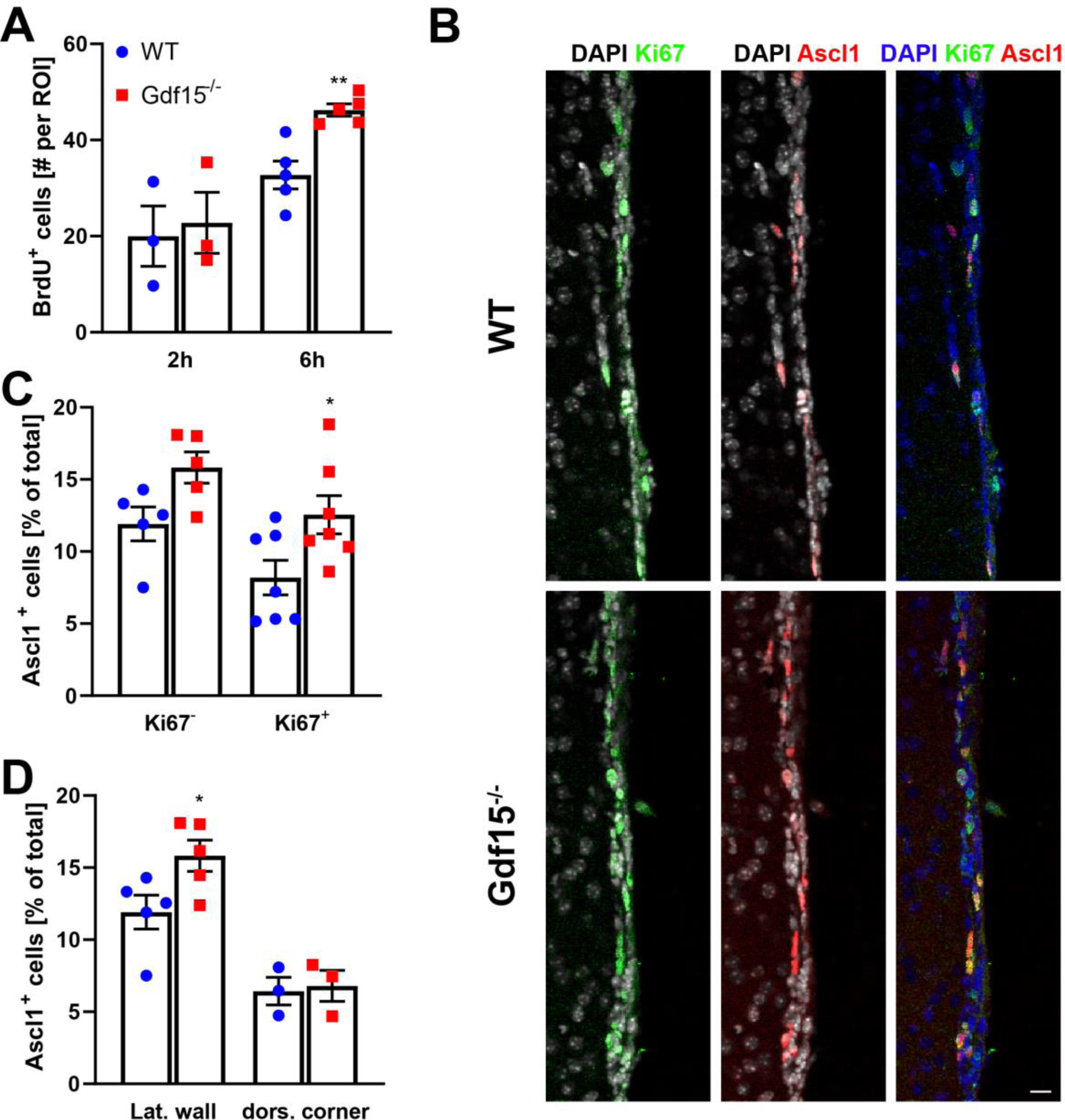
Increased proliferation in Gdf15^-/-^ animals leads to an increase in Ascl1^+^ progenitors in the lateral-ventral V/SVZ. (A) Quantitative analysis of the percentage of DAPI stained nuclei displaying BrdU immunoreactivity per region of interest (ROI) within the GE of E18 GE from WT and Gdf15^-/-^ embryos 2 and 6 hours after BrdU injection. (B) Coronal sections of the V/SVZ of adult WT and Gdf15^-/-^ mice after immunostaining for Ki67 (green) and Ascl1 (red). DAPI was used as nuclear counterstain. Scale bar = 20 µm. (C, D) Quantitative analysis of the percentage of DAPI stained nuclei in the V/SVZ of the adult WT and Gdf15^-/-^ animals immunoreactive to Ascl1 and displaying Ki67 immunoreactivity as indicated. Lat. wall = lateral wall, dors. corner = dorsolateral corner. Bars represent mean ± SEM; * indicates significance: *p<0.05, **p<0.01.

### GDF15 regulates the number of NSCs and ependymal cells

Apically dividing progenitors in the developing GE will give rise to ependymal and adult NSCs, which to a great extent derive from a common progenitor undergoing proliferation around mid-development (Ortiz-Alvarez et al., 2019) by a mechanism that involves BMP-mediated control of cell cycle regulators (Omiya et al., 2021). Moreover, both apical NSCs and ependymal cells continue to express Prominin in the postnatal SVZ (Baur et al., 2022; Carrillo-Garcia et al., 2010; Khatri et al., 2014). Therefore, we next investigated whether the increase in the proliferation of apically dividing progenitors also affects the generation of these cell types by examining the effect of the genotype on the number of multiciliated ependymal cells and progenitors displaying only one primary cilium at the apical surface of new-born mice at postnatal day 2 (P2). We here used β-catenin to label cell-cell contacts and fibroblast growth factor receptor 1 oncogene partner (FOP), a centrosomal protein, thereby visualizing cell boundaries and the basal body of the cilia, respectively. As both β-catenin and FOP-antibodies where derived from the same host species, the antigens were labelled in a single fluorescent channel and differentiated based on label localisation and intensity (Fig. 6A). Cells with a single centrosome or centrosome pair (one to two FOP^+^ dots) were counted as single-ciliated (SC), whereas cells with more than two centrosomes, i.e. multiciliated cells, were counted as ependymal (Epen; Fig. 6A). This analysis revealed a significant increase in multiciliated ependymal as well as a trend increase in SC cells in the mutant tissue compared to the WT counterpart and a significant increase in the number of total cells (Fig. 6B-D). Consistent with that, we found that in the adult V-SVZ, where multiciliated ependymal cells are fully differentiated, the number of both ependymal cells and apical NSCs, recognized on the basis of the elongated cell morphology, GFAP expression and absence of motile cilia, were increased (Fig. 6E, F). Consistent with this observation the total number of P^+^ cells, which in the adult V-SVZ include both ependyma and NSCs (Khatri et al., 2014) increased (Fig. 6G). Moreover, the incidence of clonogenic cells was not affected by the genotype (Supplementary Table S6). Taken together, these data show that total NSCs are increased in the absence of GDF15. Together, our data demonstrate that GDF15 controls the number of ependymal and NSCs in the adult V-SVZ, by regulating proliferation of apically dividing embryonic progenitors.

**Figure 6:**
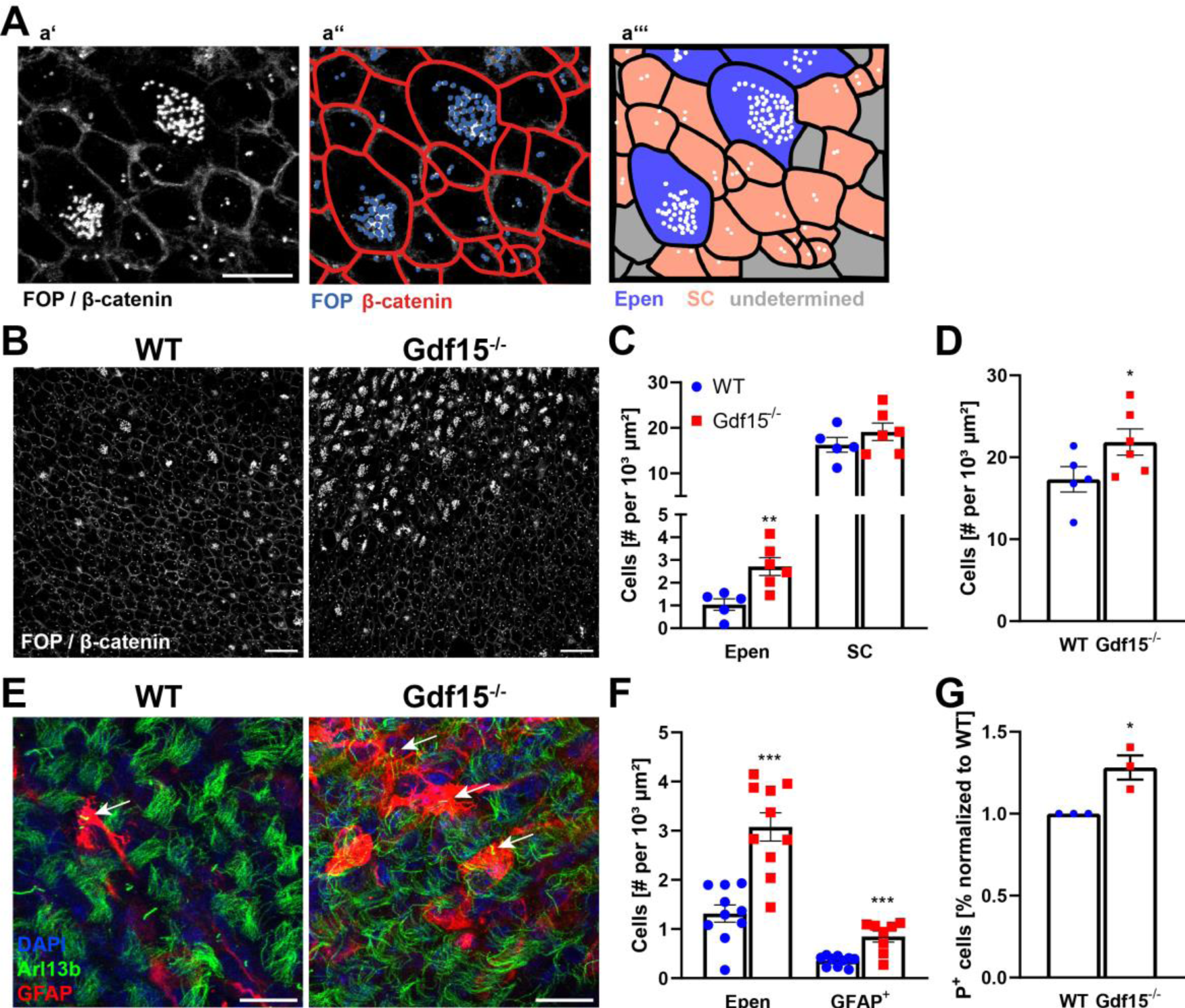
Gdf15^-/-^ animals have persistently increased number of ependymal cells and GFAP^+^ apical NSCs. (A) Schematic showing counting of ependymal (Epen) and single-ciliated (SC) cells using FOP and β-catenin as markers. (a’) Closeup of WT image in (B), showing β-catenin, indicating cell-cell-contacts, and FOP, indicating ciliary basal bodies/centrosomes, in a single channel. Scale bar = 10 µm. (a’’) β-catenin and FOP labels are distinguished by location, morphology and label intensity, with FOP being single dots that are more intense than β-catenin and located within the cell boundaries. (a’’’) Cells containing one or two centrosomes were considered SC cells (red), while cells with more than two centrosomes were considered multiciliated and therefore Epen (blue). (B) Whole mounts of the E18 lateral GE immunofluorescently labelled with FOP and β-catenin. Scale bars = 20 µm. (C, D) Quantification of cells in (B) that contain multiple basal bodies (ependymal cells, Epen) and single-ciliated cells (SC; C) as well as total cells (D). (E) Whole mounts of the adult lateral V/SVZ immunofluorescently labelled with Arl13b, to label cilia, and GFAP, to label radial glia-like cells. White arrow indicates single-ciliated NSC. Scale bars = 20 µm. (F) Quantification of cells in (E) displaying multiple cilia (Epen) or displaying GFAP immunoreactivity (GFAP^+^). (G) FACS analysis of dissociated P8 V-SVZ cells labelled with antibodies against Prominin-1 (P). Bars represent mean ± SEM; * indicates significance: *p<0.05, **p<0.01, ***p<0.001.

### Lack of GDF15 affects EGFR expression and signalling dynamics in neural progenitors

In the developing telencephalon, the acquisition of EGF responsiveness (Ciccolini, 2001) and of increase in GDF15 expression both occur at later developmental ages. Moreover, we have previously shown that GDF15 affects EGFR expression in the developing hippocampus (Carrillo-Garcia et al., 2014). Therefore, it is possible that in this region GDF15 also affects EGFR expression. Qualitative comparison of sections the GE of WT and Gdf15^-/-^ E18 embryos revealed that EGFR was more expressed at the basal than the apical side of the VZ in both the WT and Gdf15^-/-^ GE, although in the first the EGFR^+^ cells were more radially oriented than in the latter (Fig. 7A white arrows). Closer analysis of the immunostained cells, highlighted that EGFR immunoreactivity mainly displayed a punctuate appearance in the embryonic GE (Fig. 7B) and especially in the adult V-SVZ (Fig. 7C) unlike in the WT counterpart where immunoreactivity highlighted also the cell borders. Quantitative analysis also revealed a trend-decrease of fluorescence levels associated to EGFR immunoreactivity in the embryonic (Fig. 7D) and adult (Fig. 7E) tissue, which was significant only in the latter. However, measuring surface EGFR levels in apical embryonic progenitors by flow cytometry revealed a significant reduction in the number of cells displaying high levels of EGFR (E^h^) at the cell membrane in the GE, independent of Prominin expression (supplementary Fig. S6), a reduction that could be rescued by a short exposure to exogenous GDF15 (Supplementary Fig. S6A, B). Notably, the reduction in surface EGFR was observed only when EGFR was measured by binding of the less sensitive EGF-Alexa 488 (supplementary Fig. S6C, D) but not EGF-Alexa 647 (supplementary Fig. S6E, F), which unlike the first allows the detection of EGFR-expressing cells displaying also moderate levels of surface EGFR. Moreover, similar amounts of *Egfr* transcripts were expressed in the mutant and WT GE (supplementary Fig. S6G). Taken together, these data indicate that, rather than a general downregulation of EGFR, lack of GDF15 leads to a decrease in the levels of EGFR expressed at the cell membrane. We next investigated whether differences in EGFR expression are associated with changes in EGFR activation. We first measured endogenous EGFR activation by immunostaining with antibodies recognizing phosphorylated EGFR (pTyr^1092^; phEGFR). At both ages there was no significant change in endogenous EGFR activation (Fig. 7F, G). Notably, there was not a clear overlap between phEGFR and EGFR immunoreactivities, with the first being mostly expressed in weakly EGFR^+^ cells in the embryonic GE (Fig. 7B) and especially in the adult V-SVZ (Fig. 7C). The lack of correlation between EGFR expression and phosphorylation is known, since EGFR protein is degraded in the presence of strong EGFR activation (Tomas et al., 2014). Therefore, we next investigated the dynamics of EGFR activation in WT and mutant progenitors. For this, we first measured the ability of a short exposure to exogenous EGF to induce ERK phosphorylation in acutely dissociated cells from the E18 GE. This analysis revealed that independent of the genotype, both cell groups responded to EGF application (Supplementary Fig. S7A). Moreover, a short application of GDF15 was also able to induce phosphorylation of ERK, consistent with the fact that it triggers GFRAL dependent signalling. However, the dynamic response to EGFR stimulation was affected by the genotype, since levels of phERK were significantly downregulated already one hour after EGF application in mutant but not WT cells (supplementary Fig. S7B) and a lower concentration of EGF was necessary to evoke a significant response in WT than in mutant cells (Supplementary Fig. S7C). Thus, although both WT and mutant progenitors are capable to respond to EGF, the dynamics of EGFR activation differ between the two groups of progenitors, further indicating that in mutant cells the receptor is more rapidly degraded upon ligand activation resulting in overall lower EGFR levels. Supporting this hypothesis, independent of the genotype, we observed a strong colocalization between phH3 and EGFR immunoreactivity including at the apical border (Fig. 8A, B), again underscoring that the difference in EGFR between WT and mutant progenitors essentially reflect levels of EGFR expression and suggesting that also in the GE EGFR affects proliferation. Indeed, pharmacological blockade of EGFR signalling with PD158780 (PD) reduced the number of cycling progenitors (Fig. 8C, D) and dividing cells (Fig. 8E, F) in both genotypes. Although, the effect of the treatment on mitosis was significant only in the mutant GE, this likely reflects the fact that in the WT GE the significance of the effect is more difficult to quantify, due to the low number of mitotic cells already in control conditions. On the contrary, independent of the genotype, exposing the GE to exogenous EGF for 24 hours did not greatly affect the number of cycling progenitors, indicating saturation of endogenous EGFR activation. Nevertheless, the treatment caused a small but significant increase in the number of apically dividing cells in the WT tissue whereas in the mutant GE led to a decrease in the number of apically and subapically dividing progenitors that was significant only in the latter group (Fig. 8E, F). Taken together, these data show that proliferation in the mutant GE is more reliant on endogenous EGFR activation than in the WT counterpart. This is consistent with our analysis indicating that lower levels of EGFR in mutant progenitors reflect increased downregulation of EGFR expression upon activation, compared to the WT counterpart. They also show that direct manipulation of EGFR activation does not rescue the proliferation of mutant progenitors, like exposure to exogenous GDF15. A similar effect of GDF15 on EGFR expression was previously observed also in hippocampal neural progenitors, where we found that GDF15 requires active CXCR4 signalling to increase the number of EGFR^h^ hippocampal progenitors (Carrillo-Garcia et al., 2014). Indeed, in the embryonic GE blockade of CXCR4 signalling by medium application of AMD3100 (AMD) also prevented the increase in number of EGFR^h^ progenitors mediated by GDF15 (supplementary Fig. S8A) and addition of AMD to the culture medium affects specifically the ability of EGFR^h^ progenitors to undergo clone formation (supplementary Fig. S8B). Since AMD blockade affects specifically EGFR proliferating progenitors, we applied it *in situ*, to investigate how many of the proliferating EGFR^h^ cells are present in the WT and mutant GE. Treatment with AMD overnight led to a significant decrease both in terms of total cycling cells and apically dividing progenitors in the WT and mutant tissue (supplementary Fig. S8C-F). Independent of the location, no mitotic cells could be observed in AMD-treated WT tissue, whereas in the mutant tissue mitosis was significantly decreased only in apically dividing progenitors upon CXCR4 blockade (supplementary Fig. S8E, F). Taken together, these data indicate that some of the apically dividing progenitors and SAPs represent EGFR^h^ cells and that these cells, which are more reliant on active CXCR4 to undergo proliferation, are more represented in the WT GE than in the mutant counterpart. Thus, independent of GDF15 expression, endogenous EGFR signalling underlies the ability of apical GE progenitors to divide. However, whereas in the WT GE dividing more dividing progenitors are represented by EGFR^h^ cells which rely on CXCR4 activity, this population is less represented within the population of dividing progenitors in the mutant GE.

**Figure 7:**
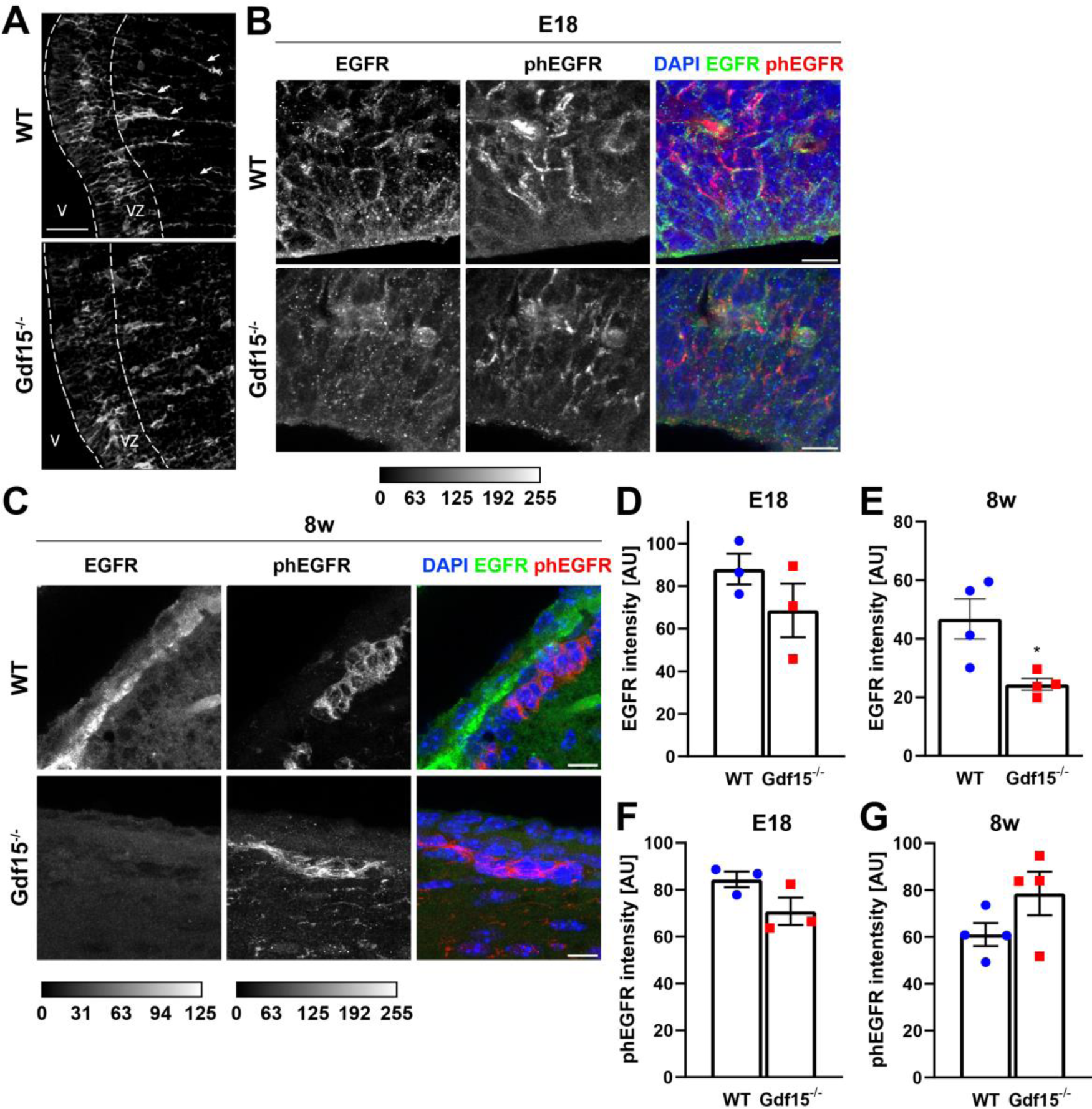
Lack of GDF15 reduces surface EGFR expression in the embryonic GE. (A) Immunofluorescent staining for EGFR in slices from E18 mice. White arrows show the columns of radially oriented EGFR labelled cells that are absent in the Gdf15^-/-^ GE. V = ventricle, VZ = ventricular zone; scale bar = 50 µm. (B) Immunofluorescent staining for EGFR and phEGFR in E18 coronal brain sections of WT and Gdf15^-/-^ mice. Scale bars = 10 µm. (C) Immunofluorescent staining for EGFR and phEGFR in 8w coronal brain sections of WT and Gdf15^-/-^ mice. Scale bars = 10 µm. (D-G) Quantification of EGFR (D, E) and phEGFR (F, G) immunofluorescence intensity of the immunostaining of E18 (D, F) and 8w animals (E, G). Bars represent mean ± SEM; * indicates significance: *p<0.05, **p<0.01.

**Figure 8:**
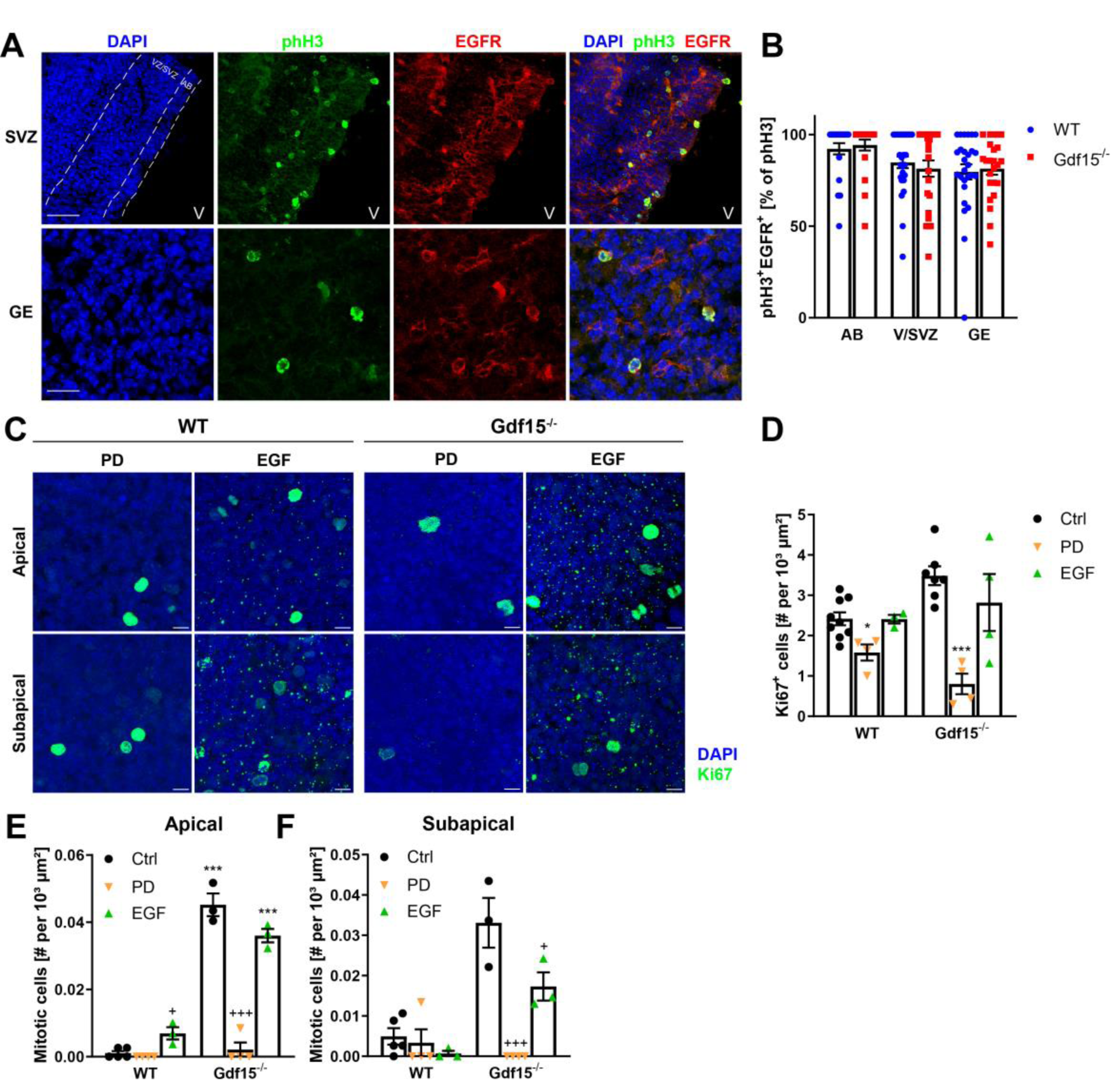
EGFR signalling is necessary for GE progenitor proliferation and is affected by lack of GDF15. (A) Coronal sections of the GE germinal region of E18 WT and Gdf15^-/-^ mice after immunostaining for phH3 (green) and EGFR (red). DAPI was used as nuclear counterstain. AB = apical border, VZ/SVZ = ventricular zone/subventricular zone, GE = deep ganglionic eminence, V = lateral ventricle. Scale bar = 37.5 µm. (B) Quantification of phH3^+^ cells co-expressing EGFR in the different parts of the niche. AB = Apical border. (C) Immunofluorescent staining of E18 GE whole mounts for proliferation marker Ki67 (green). Images are taken at the apical or subapical level of the GE as indicated. The dissected tissue was incubated in either PD158780 (PD) or recombinant human EGF for 24 hours before fixation. DAPI was used as nuclear counterstain. Scale bars = 10 µm. (D-F) Quantification of Ki67^+^ cells (D) or mitotic cells (E, F) at the apical side of E18 GE whole mounts either untreated (Ctrl) or incubated with EGFR-blocker PD or EGF as indicated. Bars represent mean ± SEM; */^+^ indicate significance: */^+^p<0.05, **/^++^p<0.01, ***/^+++^p<0.001; * indicates significance to respective WT control, ^+^ indicates significance to respective untreated control.

## Discussion

Earlier studies indicated that the neonatal periventricular zone and the choroid plexus are the main sources of GDF15 within the brain (Böttner et al., 1999; Schober et al., 2001). However, the role of GDF15 in this region remains unknown. Our data provide first evidence that GDF15 plays a role in the regulation of growth factor responsiveness, cell cycle kinetics and cell survival ultimately regulating the number of adult NSCs and ependymal cells.

It is known that most ependymal cells are born during mid late development and although becoming postmitotic they differentiate only after birth (Redmond et al., 2019; Spassky et al., 2005). Around the same time, NSCs begin a transition into quiescence, which continues into the early postnatal weeks (Borrett et al., 2020; Fuentealba et al., 2015; Furutachi et al., 2015). The generation of ependymal cells and NSCs is linked as they share a common progenitor (Ortiz-Alvarez et al., 2019). Our analysis focuses here on the effect of GDF15 on apical NSCs, which we have recently shown represent a minor pool of NSCs in the adult V-SVZ, are characterized by Prominin-1 expression and display an apical membrane and a primary cilium (Baur et al., 2022). Since basal NSCs likely derive from apical NSCs (Obernier et al., 2018), it is possible that also the number of the latter will be increased in the absence of GDF15, however this issue remains to be investigated.

The increase of GDF15 expression at late stages of development is suggestive of a developmental role for the growth factor in the regulation of the number of common progenitors and underscores a striking temporal correlation between the increase in GDF15, changes in cell cycle dynamics and in EGFR expression observed during development. This affects multiple aspects of neural progenitor behaviour including proliferation (Burrows et al., 1997; Ciccolini and Svendsen, 1998; Craig et al., 1996; Gritti et al., 1999; Suh et al., 2009), migration (Aguirre et al., 2005; Caric et al., 2001; Ciccolini et al., 2005), differentiation (Ivkovic et al., 2008), and survival by activating multiple downstream pathways (Cochard et al., 2021). Supporting a role for GDF15 in orchestrating these developmental changes, we found that short exposure to exogenous GDF15 reduces progenitor proliferation and promotes EGFR expression, showing that the phenotype observed in the mutant V-SVZ is not the result of a selective process.

At a cellular level, we found that GDF15 decreases the pool of cycling cells, apical cell division and it slows cell cycle progression. However, GDF15 is not required for progenitors to permanently exit the cell cycle and differentiate. Instead, our data *in vitro* and *in vivo* support the notion that the extra progenitor cells generated in the mutant germinal niche will proceed to differentiate in a normal manner leading to the generation of supernumerary ependymal cells and NSCs and to neuronal progenitors that will undergo compensatory cell death. Compensatory cell death was also observed upon increase of neural progenitor proliferation following loss of mCD24 expression (Belvindrah et al., 2002), suggesting that it may represent a common route of regulation of neurogenesis in the postnatal niche. However, in the mutant embryonic GE, the increase in the number of Ascl1^+^ cells was not associated with extra cell death. Since many of the cells generated at E18 in the GE migrate away and undergo terminal differentiation in the dorsal telencephalon, it is possible that the supernumerary cells generated in the Gdf15^-/-^ GE eventually undergo compensatory cell death in other regions of the telencephalon and therefore escaped our analysis. Interestingly, the increase in GDF15 expression temporally coincides with apical RG progenitors starting to withdraw from cell cycle and transitioning to the state of quiescent adult NSCs. Gene signature analysis has revealed that this transition involves shutting down several biological processes occurring between E17 and postnatal day P6 (Borrett et al., 2020), and that upon reactivation, adult NSCs reacquire characteristics of RG. Indeed, we here found that upon activation adult NSCs display the same increase in cell cycle kinetics as observed in embryonic progenitors.

Our data indicate that at the molecular level GDF15 may affect at least in part the proliferation of apical progenitors by modulating EGFR expression. Indeed, a brief exposure to exogenous GDF15 increased EGFR expression, and modulation of EGFR signalling affected the proliferation and number of apical progenitors. From late developments onwards, EGFR expression is also upregulated in NSCs and EGF is the main mitogen for NSCs (Cesetti et al., 2009; Ciccolini, 2001). Extending these observations, we here show a close association between EGFR expression and proliferation also *in vivo*, as independent of the genotype most mitotic cells expressed EGFR. Moreover, our data indicate that levels of EGFR expression affect the proliferative response to EGF, as mutant progenitors appear more dependent on endogenous EGFR signalling to undergo fast proliferation whereas activation of EGFR had opposite effects in mutant and WT GE progenitors. That EGFR signalling is an important component of the regulation of apical progenitors is consistent with the observation that in rodents, EGFR ligands are a constant component of the cerebrospinal fluid (Van Setten et al., 1999). Moreover, at late developmental stages in the rat telencephalon, TGFα is highly expressed in the choroid plexus and in the differentiated portion of GE (Kornblum et al., 1997), indicating a maximal availability of EGFR ligands at the apical and basal border of the VZ and SVZ, respectively. Consistent with the regional expression of EGFR ligands, in the postnatal brain EGFR activation is crucial for the regulation of the differentiation of ependymal cells from RG progenitors (Abdi et al., 2019). Although our data indicate that GDF15 affects the proliferation of a common ependymal/NSC progenitor and likely EGFR transduction, we did not observe a bias in differentiation, in line with the fact that EGFR impinges on multiple pathways (Cochard et al., 2021). Additionally, that modulation of EGFR signalling did not rescue the Gdf15^-/-^ phenotype in a manner similar to GDF15 application suggests that GDF15 also works through other, as of yet unknown, mechanisms in NSCs. Interestingly, embryonic human NSCs also overexpress GDF15 (Wang et al., 2010) and EGF is present in the human cerebrospinal fluid (Godard et al., 2003; Shnaper et al., 2009). Thus, our observation may have a potential relevance for the regulation of proliferation in human NSCs.

## Acknowledgements

K.B. was supported by the Interdisciplinary Center for Neuroscience (IZN) and the Landesgraduiertenförderung (LGF) of the Heidelberg University Graduate Academy. C.C.-G. was supported by contract research from the program “Adult neural stem cell” of the Baden-Württemberg Stiftung. J.S. acknowledges the Deutsche Forschungsgemeinschaft (UN 34/23-1).

## Author contributions

Conceptualization, project administration, funding acquisition: F.C.; Methodology: F.C. and J.S.; Supervision: K.B. and F.C.; Investigation: K.B., C.C.-G., Ş.Ş., M.v.H., G.H.-W. and C.M.; Formal analysis: K.B. and C.C.-G.; Visualization: K.B. and C.C.-G.; Writing – original draft: K.B. and F.C.; Writing – review and editing: K.B., C.C.-G., Ş.Ş., M.v.H. and F.C.

