## supplementary figures and tables for "Growth/differentiation factor 15 controls number of ependymal and neural stem cells in the ventricular-subventricular zone"

**Supplementary Table S1:** Antibodies and recombinant proteins used for flow cytometry.

| Antibody | Conjugate | Host | Company, Catalog # | Lot # | Concentration |
|---|---|---|---|---|---|
| Prominin-1 | APC | rat | Miltenyi Biotec, 130-102-197 | 5190517507 | 1:100 |
| Prominin-1 | PE | rat | Miltenyi Biotec, 130-102-210 | 5181126230 | 1:100 |
| Prominin-1 | BV421 | rat | BioLegend, 141213 | B255870 | 1:1000 |
| EGF | 488 | - | Invitrogen, E13345 | 1802775 | 1:1000 |
| EGF | 647 | - | Invitrogen, 35351 | 1700382 | 1:1000 |

**Supplementary Table S2:** Antibodies used for western blot. N/A = not available.

| Antibody | Conjugate | Host | Company, Catalog # | Lot # | Concentration |
|---|---|---|---|---|---|
| ERK | - | rabbit | Cell Signalling, 4695 | 8 | 1:1000 |
| phERK | - | rabbit | Cell Signalling, 4370S | 28 | 1:1000 |
| α-Tubulin | - | mouse | Sigma, T9026 | N/A | 1:10000 |
| anti-rabbit | HRP | goat | Jackson ImmunoResearch, 111-035-003 | N/A | 1:10000 |
| anti-mouse | HRP | goat | Jackson ImmunoResearch, 115-035-005 | N/A | 1:10000 |

**Supplementary Table S3:** Antibodies used for immunofluorescence. N/A = not available.

| Antibody | Host | Company, Catalog # | Lot # | Concentration |
|---|---|---|---|---|
| Adcy3 | rabbit | ThermoFisher, PA5-35382 | UL2902981 | 1:500 |
| Annexin V | rabbit | Abcam, ab14196 | N/A | 1:300 |
| Arl13b | mouse | UC Davis, 75-287<br>BioLegend, 857602 | 472-1JU-55<br>B323369 | 1:500 |
| Ascl1 (Mash1) | mouse | BD Pharmigen, 556604 | N/A | 1:100 |
| BrdU (for IdU) | mouse | Hybridoma Bank, G3G4 | 2/18/21 | 1:1000 |
| BrdU | mouse | Roche, 1296736 | N/A | 1:10 |
| β-catenin | mouse | Santa Cruz sc-7963 | G0318 | 1:100 |
| DCX | mouse | Santa Cruz, sc-271390 | D2720 | 1:200 |
| DCX | goat | Santa Cruz, | N/A | 1:200 |
| EGFR | mouse | Sigma, E2760 | N/A | 1:100 |
| EGFR, pTyr$^{1092}$ | rabbit | Abcam, ab40815 | GR111622-5 | 1:500 |
| β-galactosidase | chicken | Abcam, ab9361 | N/A | 1:10000 |
| GFAP | rabbit | Molecular probes, AA11122 | N/A | 1:1000 |
| GFRAL | sheep | Invitrogen, PA5-47769 | UH2824346A | 1:200 |
| phH3 | rabbit | EMD Millipore, 06-570 | 3689895 | 1:500 |
| Ki67 | rabbit | Abcam, 16667 | GR3313195-18 | 1:100 |
| O4 | mouse | Gift from Dr. Jaqueline Trotter | N/A | 1:100 |
| Olig2 | rabbit | Chemicon, AB9610 | N/A | 1:200 |
| β-Tubulin III (TuJ1) | mouse | Sigma, T8660 | N/A | 1:400 |

**Supplementary Table S4:** Number of pycnotic cells (as percentage of all cells) from the dissociated SVZ of adult WT and *Gdf15*$^{-/-}$ mice after 2 days of differentiation.

| *2 days differentiation* | Control | + GDF15 | *p*-Value | N |
|---|---|---|---|---|
| WT | 4.46 ± 0.74 | 4.04 ± 0.48 | 0.91 | 7 |
| Gdf15$^{-/-}$ | 6.19 ± 1.32 | 4.33 ± 0.46 | 0.24 | 6 |
| *p*-Value | 0.26 | 0.96 | | |

**Supplementary Table S5:** Number of pycnotic cells (as percentage of all cells) from the dissociated SVZ of adult WT and *Gdf15*$^{-/-}$ mice after 4 days of differentiation.

| *4 days differentiation* | Control | + GDF15 | *p*-Value | N |
|---|---|---|---|---|
| WT | 4.83 ± 0.37 | 4.15 ± 0.24 | 0.2 | 7 |
| Gdf15$^{-/-}$ | 6.99 ± 0.15 | 4.92 ± 0.33 | 0.0002 | 6 |
| *p*-Value | < 0.0001 | 0.15 | | |

**Supplementary Table S6:** Number of clonogenic cells (as percentage of plated) within the indicated population sorted from the dissociated SVZ of adult WT and *Gdf15*$^{-/-}$ mice.

| Clones DIV 0 | WT | *Gdf15*$^{-/-}$ |
|---|---|---|
| $P^-E^h$ cells | 14.37±7.4 | 12.71±3.9 |
| $P^+E^h$ cells | 21.3 ± 1.79 | 38.35 ± 9.51 |

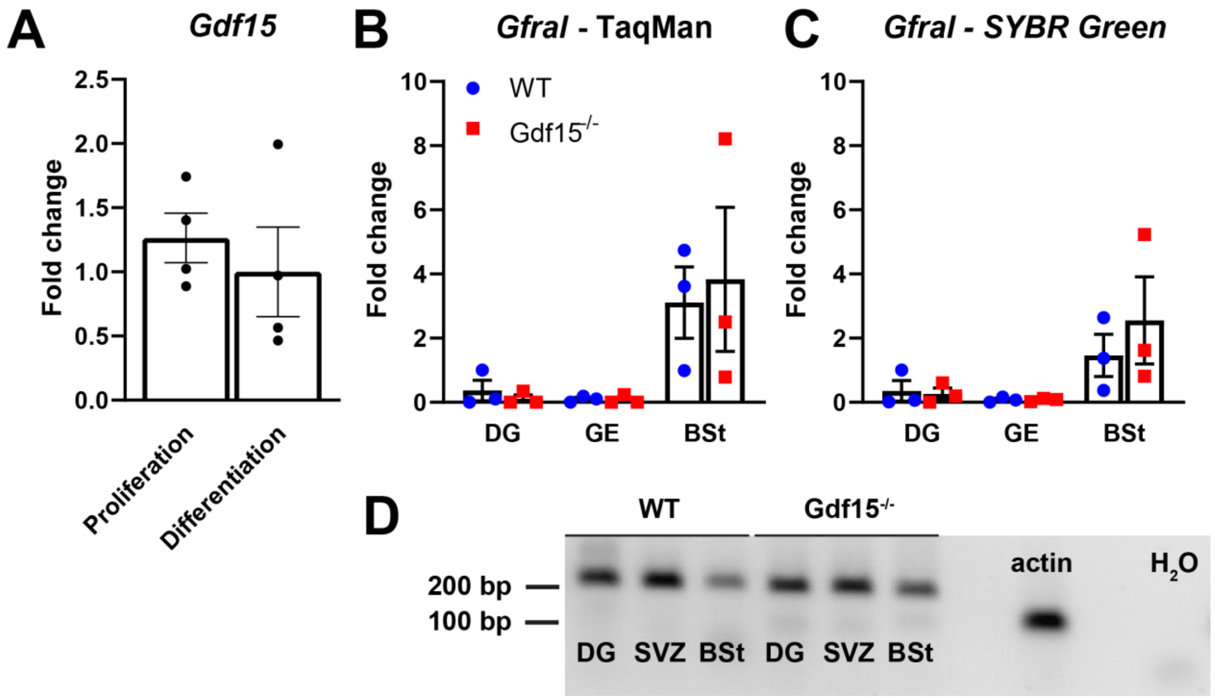


**Supplementary Figure S1: Expression of Gdf15 and Gfral.**

(A) Fold change of *Gdf15* mRNA levels during proliferation and differentiation.

(B) Fold change of *Gfral* mRNA levels in different brain regions, determined by TaqMan qPCR. Some values for DG and SVZ are 0 (undetermined).

(C) Fold change of *Gfral* mRNA levels in different brain regions, determined by SYBR Green qPCR. While expression in the DG and SVZ is markedly lower than in the BSt, it is never 0.

(D) Agarose gel with PCR products from SYBR green qPCR (C), with actin as positive control, and *Gfral* primers with $H_2O$ instead of cDNA as negative control. The PCR product for Gfral has an expected length of 213 bp and is present in all brain regions, but not in the water control.

Bars represent mean ± SEM; DG = dentate gyrus, SVZ = subventricular zone, BSt = brain stem.

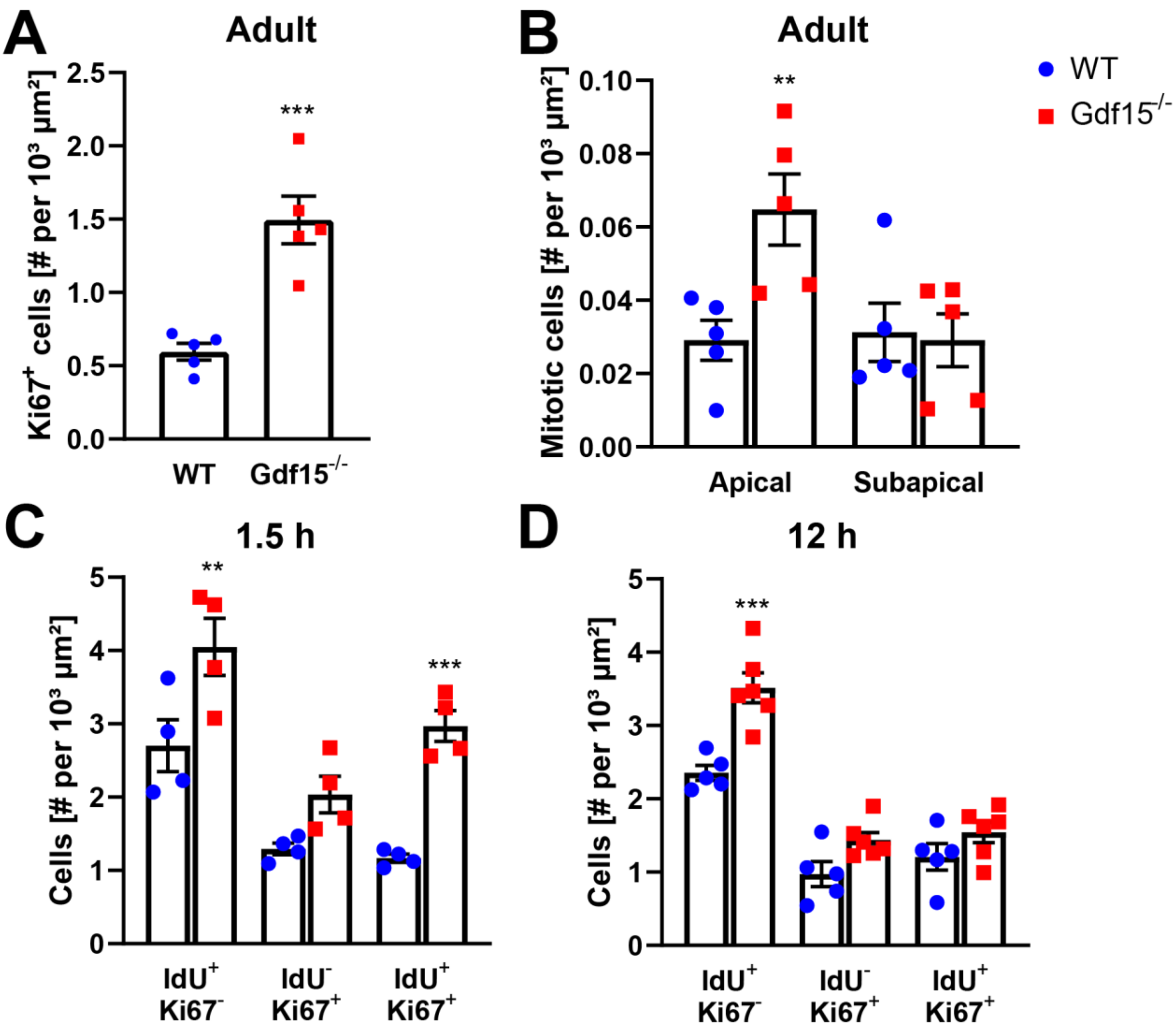


**Supplementary Figure S2: Lack of Gdf15 increases proliferation.**

(A, B) Quantification of cells in adult V/SVZ whole mounts expressing Ki67 (A) or visibly in M-phase (B).

(C, D) Quantification of cells positive for IdU and/or Ki67 immediately (C) or 12 hours after IdU application (D).

Bars and data points represent mean ± SEM; * indicates significance compared to WT control: **p<0.01, ***p<0.001.

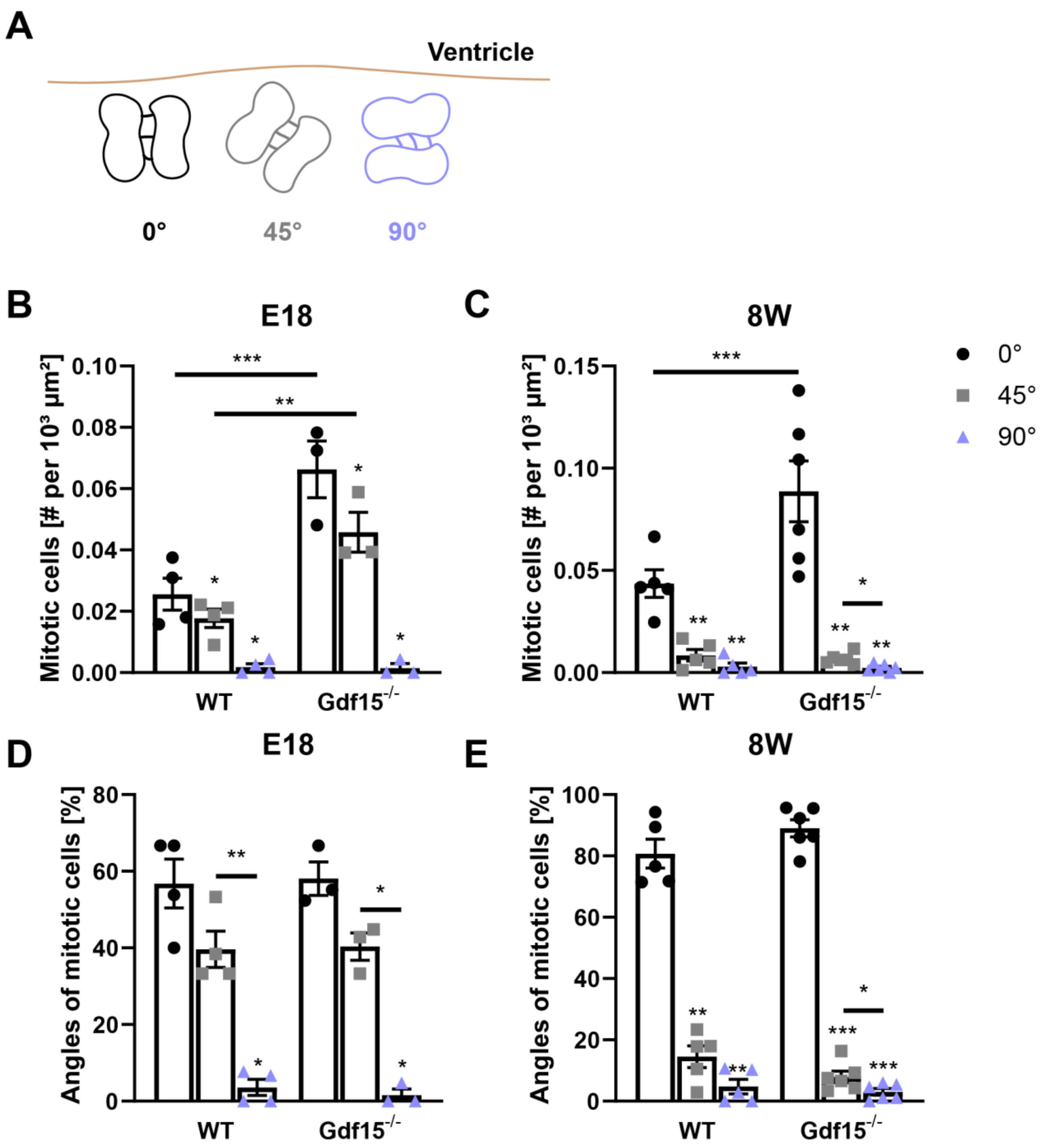


**Supplementary Figure S3: GDF15 KO does not change mode of division.**

(A) Sketch illustrating the different division modes determined by the angle of the mitotic spindle relative to an orthogonal axis to the ventricular surface. Categories were determined as follows: 0° = 0°-30°; 45° = 30°-60°; 90° = 60°-90°.

(B, C) Quantification of total number of mitotic cells according to angle of mitosis at E18 (B) and in adult animals (C).

(D, E) Quantification of mitotic cells according to angle normalized to total mitotic cells at E18 (D) and in adult animals (E).

Bars represent mean ± SEM; * indicates significance to 0° group (on top of bars) or to other group (line): *p<0.05, **p<0.01, ***p<0.001.

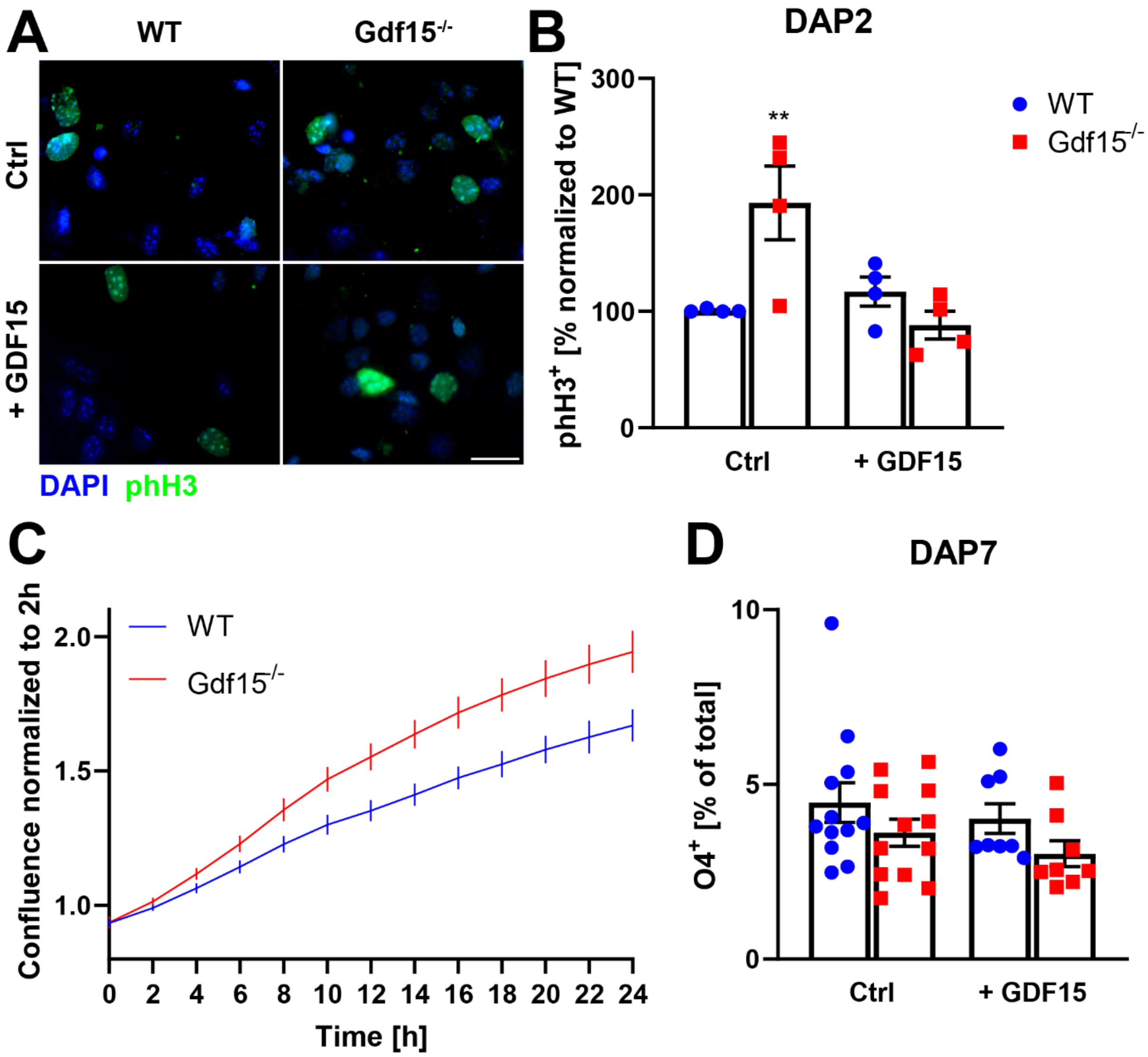


**Supplementary Figure S4: Gdf15$^{-/-}$ cells show increased proliferation *in vitro*.**

(A) Fluorescent micrographs of neutrosphere derived progenitors obtained from the GE of E18 embryos of the given genotypes. Cultures were fixed two days after plating (DAP2) in differentiating conditions in the presence of 10 ng/ml GDF15 as indicated, and immunostained with mitosis marker phH3. Scale bar = 20 µm.

(B) Quantification of phH3$^{+}$ cells (Ctrl). ** indicate significance p<0.01.

(C) Fold change of confluence of primary WT and Gdf15$^{-/-}$ cells measured by IncuCyte.

(D) Quantification of cells expressing oligodendrocyte marker O4 at DAP7.

Bars and lines represent mean ± SEM.

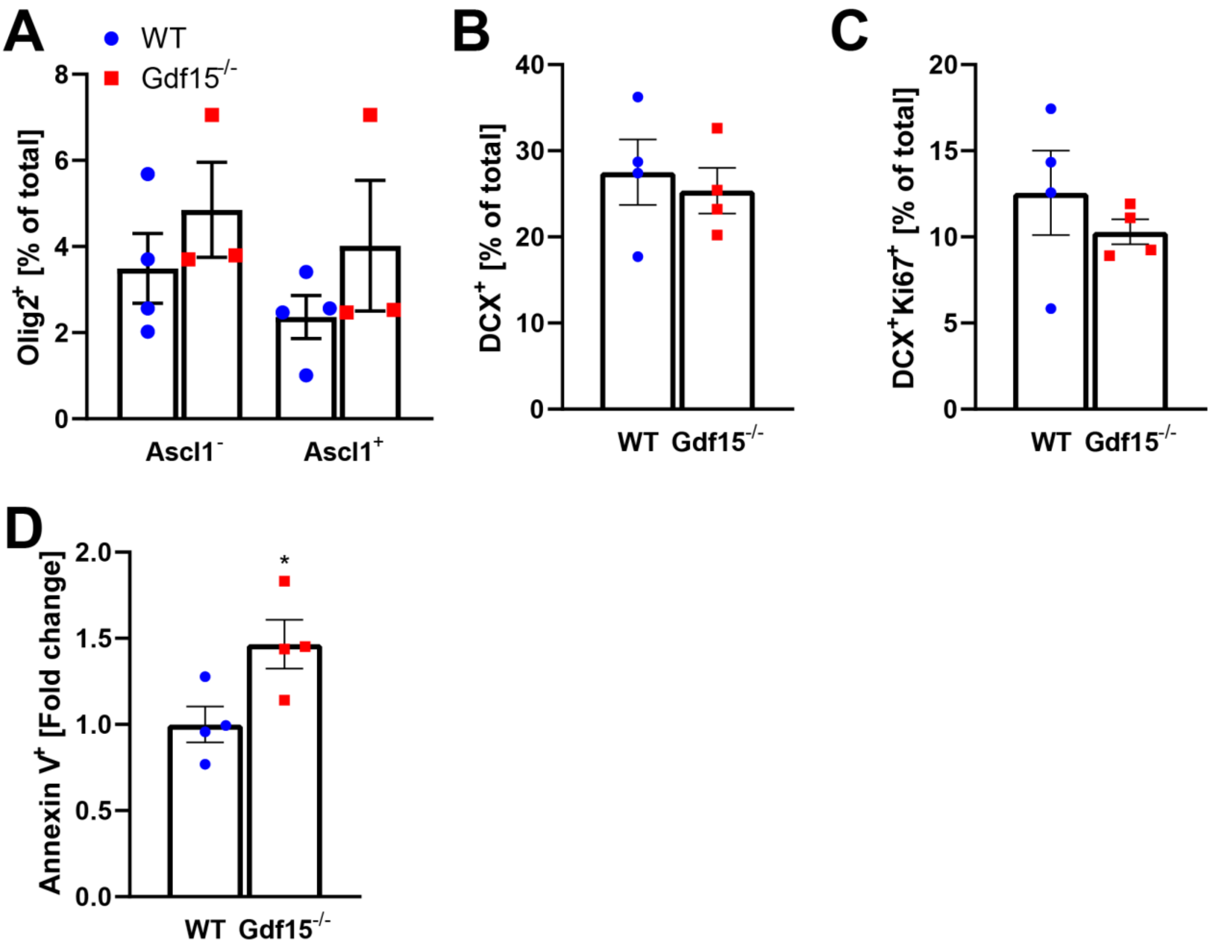


**Supplementary Figure S5: Increased proliferation does not lead to higher levels of neurons but to increased cell death.**

(A-D) Quantification of cells in adult coronal brain sections expressing oligodendrocyte marker Olig2 and neural precursor marker Ascl1 as indicated (A), neuroblast marker DCX (B, C), proliferation marker Ki67 (C) and cell death marker Annexin V (D). Bars represent mean ± SEM; * indicates significance $p<0.05$.

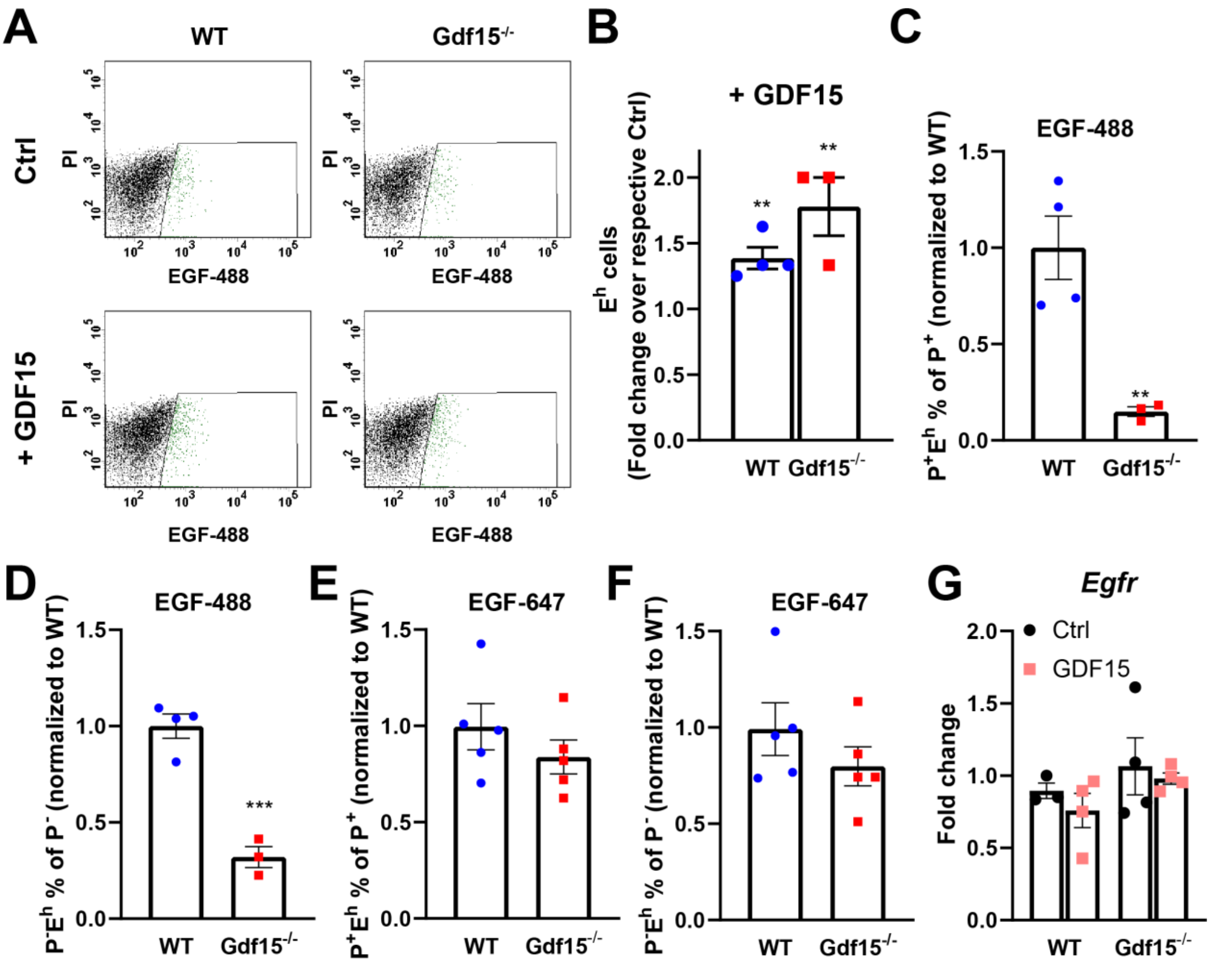


**Supplementary Figure S6: Gdf15$^{-/-}$ progenitors in the E18 GE have lower levels of surface EGFR but not overall lower expression.**

(A) Flow cytometry scatter plots of cells derived from the E18 WT and Gdf15$^{-/-}$ GE with and without GDF15 treatment (+ GDF15 and Ctrl, respectively), showing levels of surface EGFR expression after tagging with immunofluorescently labelled EGF ($E^h$, green dots).

(B) Fold change in EGFR-expressing ($E^h$) cells isolated from the WT and Gdf15$^{-/-}$ GE at E18 as analysed by flow cytometry. Cells were incubated in medium without or with GDF15 as indicated, and the percentage was normalized to the untreated control.

(C-F) Surface expression of EGFR in relation to Prominin-1-expression as determined by FACS, normalized to WT levels, using less sensitive EGF-488 (C, D) or more sensitive EGF-647 (E, F).

(G) *Egfr* mRNA transcript levels in WT and Gdf15$^{-/-}$ SVZs, determined by qPCR. 24 h before sample harvest, whole SVZ extracts were incubated in medium without (Ctrl) and with GDF15.

Bars represent mean ± SEM; * indicates significance: **$p<0.01$, ***$p<0.001$.

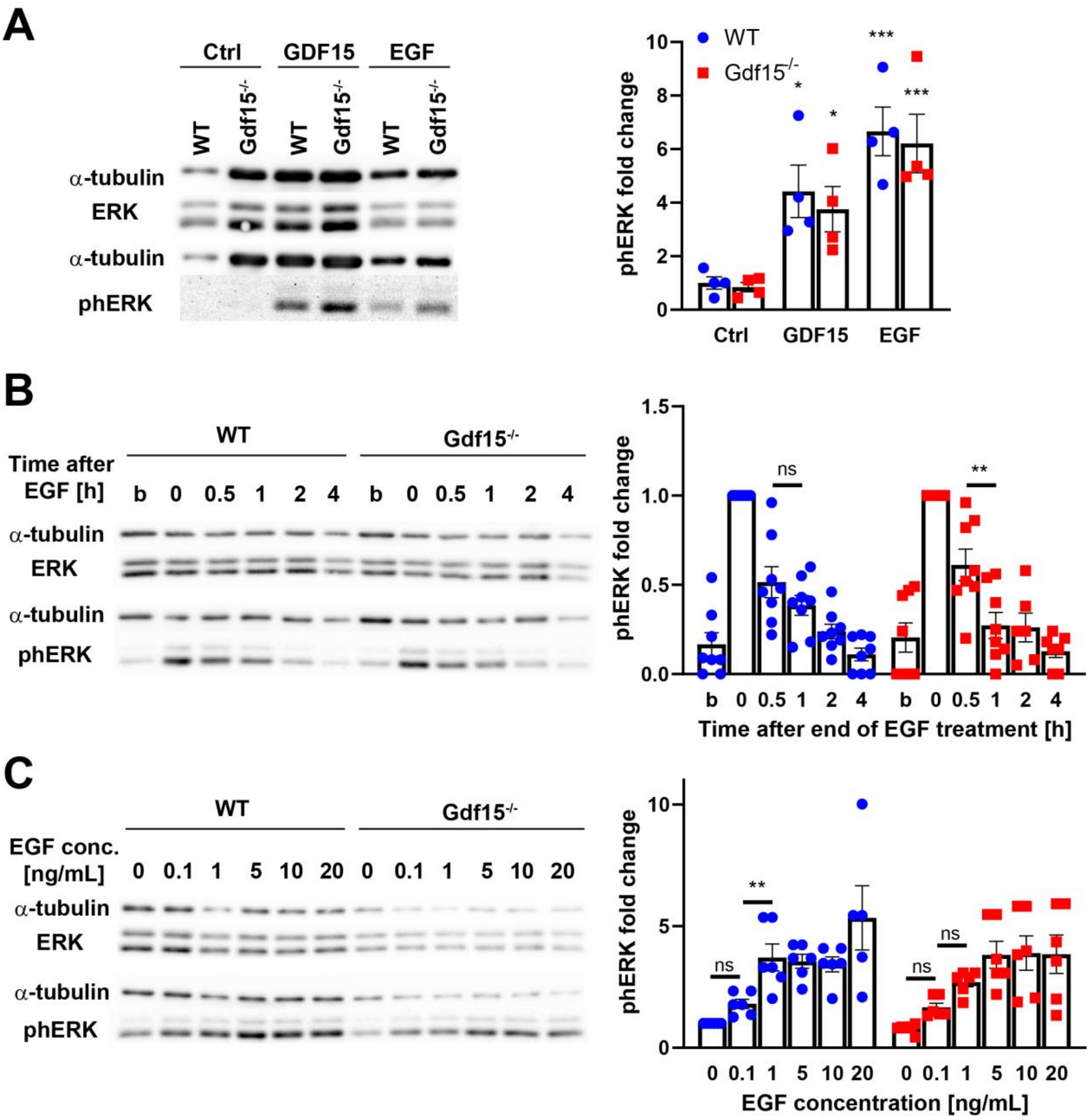


**Supplementary Figure S7: WT and Gdf15$^{-/-}$ show differences in EGFR activation kinetics.**

(A) Western blot and quantification of phERK levels in cells incubated with GDF15 for 24h or EGF for 7 minutes.

(B) Western blot and quantification of phERK levels in WT and Gdf15$^{-/-}$ cells incubated without (baseline, b) or with EGF for 7 minutes and then further incubated without EGF for 0, 0.5, 1, 2 or 4 hours as indicated.

(C) Western blot and quantification of phERK levels in WT and Gdf15$^{-/-}$ cells incubated with 0, 0.1, 1, 5, 10 or 20 ng/ml EGF for 7 minutes as indicated.

Bars represent mean ± SEM; ns = not significant; * indicates significance: *p<0.05, **p<0.01, ***p<0.001.

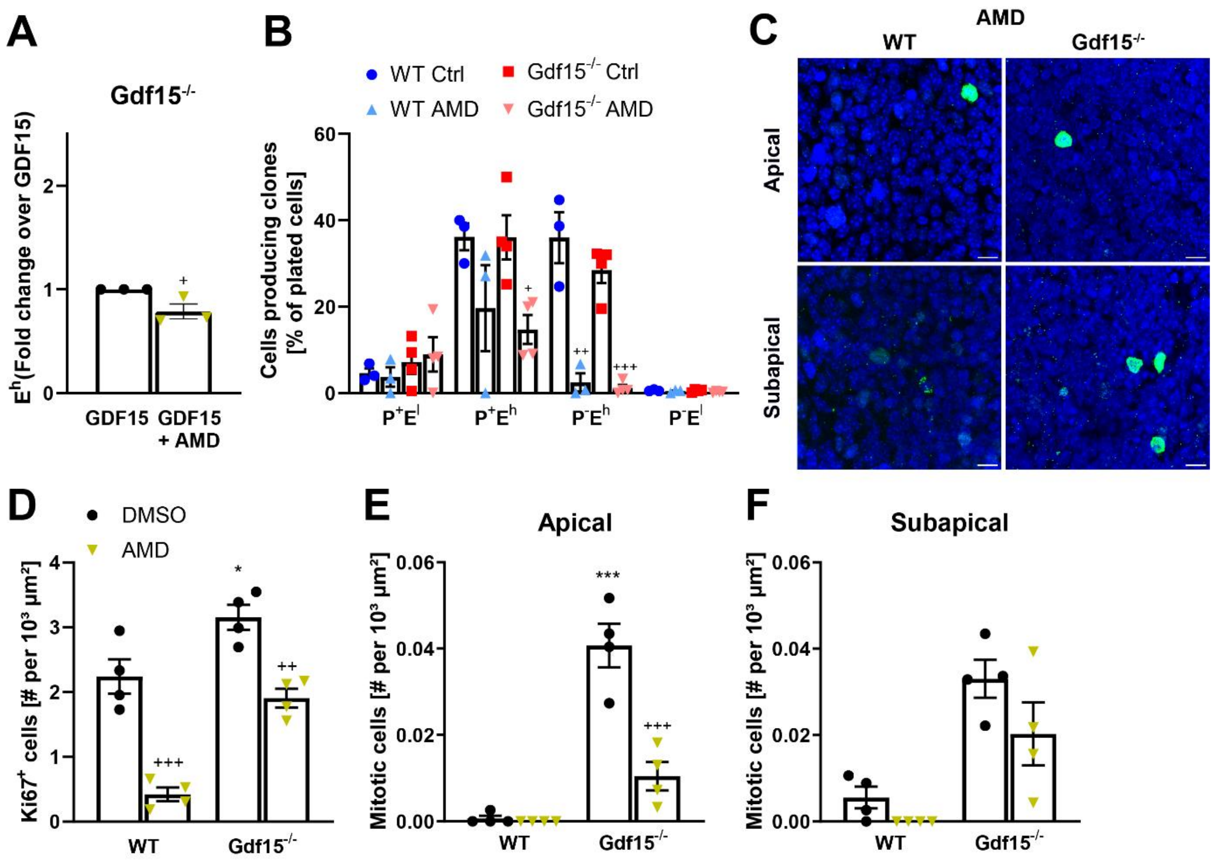


**Supplementary Figure S8: Effects of CXCR4 inhibition.**

(A) Fold change in EGFR-expressing ($E^h$) cells isolated from the Gdf15$^{-/-}$ GE at E18 as analysed by flow cytometry. Cells were incubated in medium with GDF15, without or with addition of CXCR4 antagonist AMD3100 (AMD).

(B) Clonal analysis of cells treated with AMD3100 after FACS sort.

(C) Immunofluorescent staining of E18 GE whole mounts for proliferation marker Ki67 (green). Images are taken at the apical or subapical level of the GE as indicated. The dissected tissue was incubated with 6 µM AMD3100 for 24 hours before fixation. DAPI was used as nuclear counterstain. Scale bars = 10 µm.

(D-F) Quantification of Ki67$^+$ cells (D) or dividing cells (E,F) at the apical side of E18 GE whole mounts either incubated with or with DMSO as control or CXCR4-blocker AMD overnight.

Bars represent mean ± SEM; $^+$ indicate significance to Control: $^+p<0.05$, $^{++}p<0.01$, $^{+++}p<0.001$.
